# GRaNIE and GRaNPA: Inference and evaluation of enhancer-mediated gene regulatory networks applied to study macrophages

**DOI:** 10.1101/2021.12.18.473290

**Authors:** Aryan Kamal, Christian Arnold, Annique Claringbould, Rim Moussa, Nila H. Servaas, Maksim Kholmatov, Neha Daga, Daria Nogina, Sophia Mueller-Dott, Armando Reyes-Palomares, Giovanni Palla, Olga Sigalova, Daria Bunina, Caroline Pabst, Judith B. Zaugg

## Abstract

Among the biggest challenges in the post-GWAS (genome-wide association studies) era is the interpretation of disease-associated genetic variants in non-coding genomic regions. Enhancers have emerged as key players in mediating the effect of genetic variants on complex traits and diseases. Their activity is regulated by a combination of transcription factors (TFs), epigenetic changes and genetic variants. Several approaches exist to link enhancers to their target genes, and others that infer TF-gene connections. However, we currently lack a framework that systematically integrates enhancers into TF-gene regulatory networks. Furthermore, we lack an unbiased way of assessing whether inferred regulatory interactions are biologically meaningful. Here we present two methods, implemented as user-friendly R packages: GRaNIE (Gene Regulatory Network Inference including Enhancers) for building enhancer-based gene regulatory networks (eGRNs) and GRaNPA (Gene Regulatory Network Performance Analysis) for evaluating GRNs. GRaNIE jointly infers TF-enhancer, enhancer-gene and TF-gene interactions by integrating open chromatin data such as ATAC-Seq or H3K27ac with RNA-seq across a set of samples (e.g. individuals), and optionally also Hi-C data. GRaNPA is a general framework for evaluating the biological relevance of TF-gene GRNs by assessing their performance for predicting cell-type specific differential expression. We demonstrate the power of our tool-suite by investigating gene regulatory mechanisms in macrophages that underlie their response to infection and cancer, their involvement in common genetic diseases including autoimmune diseases, and identify the TF PURA as putative regulator of pro-inflammatory macrophage polarisation.

**Availability:**

- GRaNIE: https://bioconductor.org/packages/release/bioc/html/GRaNIE.html
- GRaNPA: https://git.embl.de/grp-zaugg/GRaNPA

**Graphical abstract:** 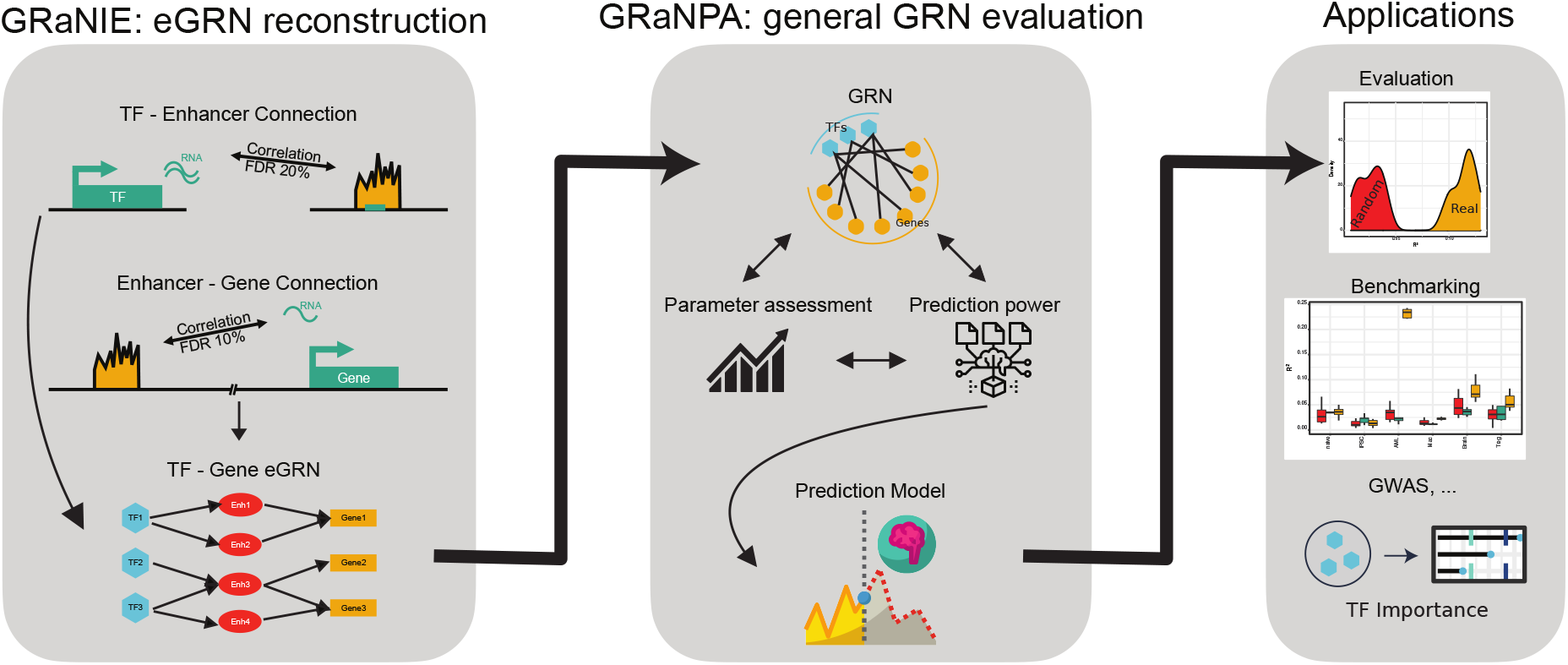

## Introduction

Enhancers are genomic locations that play an important role in cell-type specific gene regulation, and an increasing number of diseases are linked to impaired enhancer function (1,2). This is particularly obvious for common genetic traits and diseases for which genome-wide association studies (GWAS) have linked over 200,000 genetic variants with over 40,000 traits and diseases, yet the vast majority of the disease-associated genetic variants lie in non-coding regions far from promoters (1), thus likely affecting enhancers and having a regulatory role.

Among the biggest challenges in the post-GWAS era is the interpretation of these disease-associated genetic variants in non-coding genomic regions. For most disease-associated genetic variants in regulatory regions it is still unclear what genes they target. This is further complicated by the cell- and condition-specific activity of gene regulatory elements, which is likely conferred by transcription factors (TFs) that regulate the specific elements. On the one hand, a recent study points at the importance of studying TFs for understanding genetic variants associated with auto-immune diseases (3), which is corroborated by the fact that *trans*-expression quantitative trait loci (eQTLs), which are likely to act via TFs, tend to be more enriched in disease-associated variants than c/s-eQTLs (4). On the other hand, it is crucial to include putative enhancers for predicting the function of TFs in cell-fate determination (5,6). Enhancer-mediated gene regulatory networks reconstructed from single cell RNA and ATAC-seq profiling in the fly brain have led to a better understanding of the regulatory diversity across the different neuronal cell types (6). We have previously shown the power of using enhancer-based analyses to understand disease mechanisms in pulmonary arterial hypertension (7). Thus, for interpreting disease-associated genetic variants, or enhancers in general, it will be crucial to jointly investigate TF activity, enhancers and gene expression in a cell-type specific manner.

Several approaches have been proposed to infer bi-partite TF-gene networks for example based on co-expression in bulk (8,9), or single cell data (10–12), based on partial information decomposition (13), time-course data (14), or data curation (15–18). At the same time, methods for inferring enhancer-gene links have been proposed e.g. using co-variation of peaks (19–21), or targeted perturbations of enhancers followed by sequencing (19,20). However, only few approaches jointly infer TF-enhancer and enhancer-gene links (22), and for none of them a computational tool is currently available. Thus, we currently lack tools to infer networks that enable the study of context-specific interaction between TFs, regulatory elements, and genes.

An important step in regulatory network reconstruction is to evaluate their biological significance. Common approaches for assessing regulatory interactions include benchmarking them against networks from simulated data or against known biological networks (23–25). The drawback of simulated networks is that they are based on many assumptions of the structure of a “true” biological network. Benchmarking against true biological networks, while very important for assessing known connections, typically suffers from a strong literature bias (26), low complexity of these “true” circuits and a limited range of connections and cell types, and are thus not well-suited for an unbiased evaluation of GRNs. In general, each network inference method will have its own bias and shortcomings, and performance will depend on the benchmarking dataset (23–25). Thus, there is a need for an unbiased approach to assess the biological relevance of inferred and curated regulatory interactions as well as the regulon of individual TFs.

Here we present a tool-suite for building and evaluating enhancer-based gene regulatory networks (eGRNs) called GRaNIE (Gene Regulatory Network Inference including Enhancers - https://bioconductor.org/packages/release/bioc/html/GRaNIE.html) and GRaNPA (Gene Regulatory Network Performance Analysis - https://git.embl.de/grp-zaugg/GRaNPA), respectively. GRaNIE jointly infers TF-enhancer and enhancer-gene interactions based on covariation of bulk RNA-seq expression and chromatin accessibility (ATAC-seq) or ChIP-seq for active histone marks (e.g. H3K27ac) across biosamples. GRaNPA assesses the biological relevance - of any GRN - using a machine learning framework and identifies TFs that predict cell-type specific expression response to perturbations. We demonstrate that GRaNIE infers biologically meaningful eGRNs in macrophages, using ChIP-seq for validating TF-enhancer links and eQTL data for validating the enhancer-genes links. We further demonstrate the cell-type specific nature of GRaNIE-inferred eGRNs for macrophages, T-cells and AML based on cell-type specific GRaNPA evaluation and prediction of cell-type specific TF knock-out (KO) data in macrophages. Using GRaNIE followed by GRaNPA, we identify PURA as putative TF driving the pro-inflammatory polarisation of macrophages, which we corroborate with orthogonal phosphoproteomics data, and we confirm earlier observations from mice that MBD2 drives the anti-inflammatory program in macrophages. Furthermore, we find enhancers in the macrophage eGRNs enriched for autoimmune disease variants, which GRaNIE links to upstream TFs and putative target genes. Finally, we provide a comprehensive resource of cell-type specific GRNs for three other cell types (https://apps.embl.de/grn/).

## Results

### Overview and conceptual description of the GRaNIE algorithm

To interpret genetic and epigenetic variation in regulatory (enhancer and promoter) regions, here defined by ATAC-seq peaks and hereafter referred to as “peaks”, we developed GRaNIE, an R/Bioconductor package for jointly inferring TF-enhancer/promoter and enhancer/promoter-gene interactions from the same data in a context-specific manner. Conceptually, the software is based on an approach we have devised for a recent study in which we investigated enhancer-mediated disease mechanisms of pulmonary arterial hypertension (7). Briefly, GRaNIE first identifies TF-peak and peak-gene links, and then integrates them into an eGRN.

The TF-peak links are based on statistically significant co-variation of TF expression and peak accessibility across samples (e.g. individuals, recommended minimum number ~10-15), taking into account predicted TF binding sites. To obtain them, GRaNIE calculates all pairwise correlations between TF expression levels (RNA-seq) and peak signal (ATAC/ChIP-seq), stratified by whether or not the peak contains a putative binding site of the TF. For each TF it then uses the distribution of all peaks that do not contain its putative binding site as background to calculate an empirical false discovery rate (FDR) for assessing the significance of TF-peak links **(Fig. S1).** In our previous work we have demonstrated that negative TF-peak correlations indicate the TF acts as transcriptional repressor while positive correlations indicate an activator role, thus allowing the classification of TFs into activators and repressors (27). As a quality control (QC) in GRaNIE we recommend comparing the number of real TF-peak links to those obtained from an eGRN inferred from randomized data. This is implemented in GRaNIE by permuting sample labels, peak labels and motif labels and inferring an eGRN from that (run automatically in the standard workflow).

Similarly, peak-gene links are based on significant co-variation of peak accessibility and gene expression across samples **(Fig. 1A, Fig. S1).** For this, GRaNIE calculates the correlation between the expression of a gene and peak signal of all peaks within an adjustable, defined distance of its transcription start site (TSS, default is 250kb). GRaNIE also allows the use of chromatin conformation data, such as Hi-C, to define which peak-gene pairs will be tested within a 3D proximity **(Fig. S1).** Since chromatin accessibility in regulatory elements is generally associated with active gene regulation and transcription, we only expect positive correlations for functional peak-gene links. Notably, the same is still true even for repressor-bound elements, where binding of a repressor leads to loss of both accessibility and transcription (27). Negative correlations have no clear biological meaning and therefore may indicate remaining batch effects or random noise. We therefore implemented the assessment of positive versus negative peak-gene correlations as a QC metric in GRaNIE to judge the signal-to-noise ratio in the data, and by default only retain positive enhancer-gene links. In addition, comparing these QC metrics with permuted data is recommended (implemented in GRaNIE, see above).

**Figure 1:**
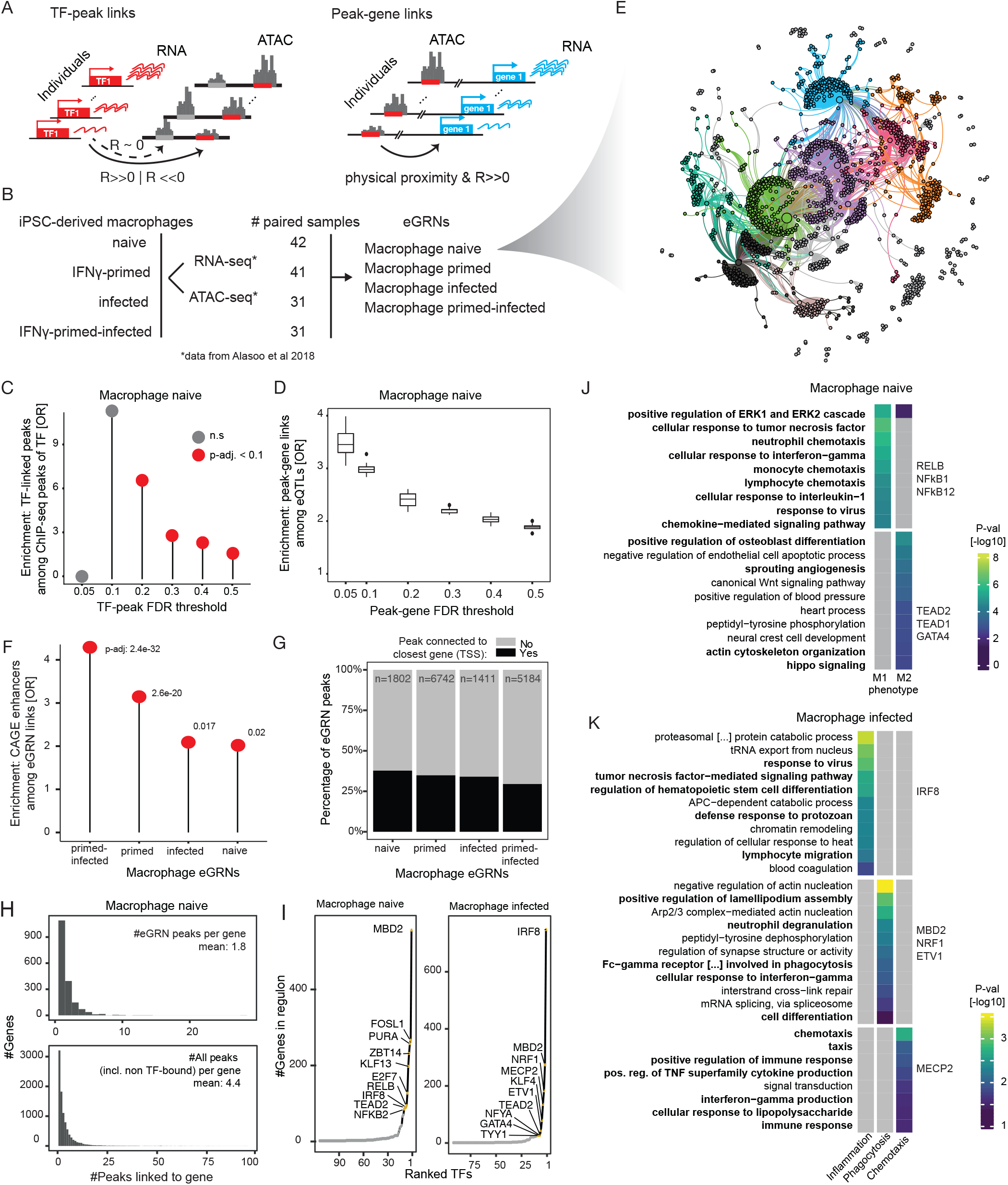
Overview, application and validation of GRaNIE. **A)** Schematic of the eGRN construction by GRaNIE, including the TF to peak (left) and peak to gene (right) links (detailed workflow in **Fig. S1). B)** Datasets used for macrophage eGRN construction and evaluation. C) Validation of the eGRN TF-peak links with ChIP-seq data. Enrichment of ChIP-seq peaks overlapping a GRaNIE-inferred TF-bound peak (same TF) are shown for different TF-peak FDRs in the naive macrophage eGRN. Background: peaks that contain the motif for the respective TF but are not significantly linked (see **Fig. S2** for other eGRNs). **D)** Validation of the eGRN peak-gene links with macrophage eQTLs. Plots show the enrichment of eGRN links overlapping an eQTL over randomly sampled distance-matched peak-gene links for different peak-gene FDRs in the naive macrophage eGRN (see **Fig. S3** for other eGRNs). **E)** Forced-directed visualization of the naive macrophage eGRN (see **Fig. S4** for the other eGRNs). The colors correspond to the identified communities. F) Enrichment of macrophage-specific FANTOM5 CAGE enhancers among the macrophage eGRN peaks. **G)** Fraction of eGRN peaks connected to the closest gene (black) vs other (grey) genes for the macrophage eGRNs. H) Number of peaks linked to a gene shown as histogram for eGRN peaks (top) and all peaks (including non-TF bound; bottom) for the naive macrophage data (see **Fig. S7** for other eGRNs). Mean number of peaks indicated in the panels. I) Number of genes connected to each TF for the naive macrophage eGRN (top 10 TFs are labelled). **J-K)** GO enrichment for selected communities from the naive (J) and infected (K) macrophage eGRN (see **Table S7** for the full table of enrichments across communities for all macrophage eGRNs).

GRaNIE then combines these two types of edges to create a tripartite TF-enhancer-gene eGRN, based on three user-defined thresholds: false discovery rate (FDR) of TF-enhancer edges, FDR of enhancer-gene edges, and maximum distance between enhancer and transcription start site (TSS).

The whole framework is implemented in a user-friendly R/Bioconductor-package. In addition to the eGRN reconstruction function, GRaNIE comprises a set of easy-to-use functions for generating and visualising network statistics, identifying network communities, performing subnetwork-specific Gene Ontology (GO) enrichment analysis, and various QC plots. An extensive documentation is available at (https://grp-zaugg.embl-community.io/GRaNIE).

### Application of GRaNIE for generating cell-type specific eGRN in macrophages

Macrophages are large white blood cells of the innate immune system that can be found in essentially all tissues and play a role in inflammatory disorders. Genetic variants associated with several auto-immune and other diseases are enriched for enhancers active in macrophages (28,29). Given that inflammatory conditions are underlying many common diseases, macrophages present an important cell type within which disease-associated variants manifest their effect. Thus, macrophages present an ideal cell type for applying and testing the eGRN framework.

We obtained joint RNA- and ATAC-seq data for (induced pluripotent stem cell) iPSC-derived macrophages from 31-45 individuals in four conditions (naive, interferon gamma (IFN-γ)-primed, infected (with Salmonella), and IFN-γ-primed-infected) (28) **(Fig. 1B).** For each of these, we used GRaNIE using TF binding sites predictions based on HOCOMOCO v11 and PWMScan as described in (27).

To assess the GRaNIE inferred eGRNs and to define reasonable default values for the two TF-peak and peak-gene FDR thresholds, we sought to obtain independent molecular evidence. Since molecular ground truth data for TF-peak-gene links doesn’t exist, we decided to evaluate both types of links independently using cell-type specific ChIP-seq data for the TF-peak links and cell-type specific eQTL data for the peak-gene links. Specifically, we obtained macrophage-specific ChIP-seq data from ReMap 2022 (Hammal et al. 2022), and quantified the enrichment of GRaNIE-inferred TF-bound peaks among ChIP-seq peaks using ATAC-peaks that contain the TF motif as background (see Methods). For the naive, the primed and the infected macrophage eGRNs, this revealed a significant enrichment for ChIP-seq signal for FDR 0.2, which steadily decreased with increasing FDR **(Fig. 1C, Fig. S2).** The primed-infected eGRN did not show any significant enrichment. For the gene-peak links we used macrophage-specific c/s-eQTLs to assess the enrichment of eQTL links in the GRaNIE links over a distance-matched set of control links at various FDRs. This revealed a steady decrease of the odds ratio with increasing FDR for all four eGRNs **(Fig. 1D, Fig. S3).** Based on these results, we chose 0.2 as default for TF-peak FDR and 0.1 as default for peak-gene FDR and we excluded the primed-infected eGRN from any further analysis.

Using these default parameters we obtained an eGRN for each of the three conditions comprising between 92 to 126 TFs, 1411 to 6742 enhancers, 1454 to 3869 genes **(Fig. 1E, Fig. S4A-C; Tables 1 and S1-S3).** For all eGRNs we observed much fewer significant connections when running GRaNIE on data with permuted sample labels, peak labels and motif labels **(Fig. S5A-D),** signifying that they pass the QC for TF-peak links. Similarly, for the peak-gene links, we find that all eGRNs show more signal for positive (expected signal) than for negative (noise) correlations, and that the signal-to-noise ratio decreases with peak-gene distance until no signal is left for random peak-gene pairs **(Fig. S6A-D).** In addition, we observed a significant enrichment for the TF-peak-gene links for being cell-type specific active enhancers based on CAGE data (30) **(Fig. 1F),** further corroborating that GRaNIE infers biologically meaningful eGRNs.

**Table 1:** Summary of the eGRNs described in the main text.

Network statistics for the various eGRNs based on default parameters (TF-peak FDR=0.2, peak-gene FDR=0.1, peak-gene distance $\leq 250$ kb, activator/repressor stringency threshold based on the 10th percentile)
| *eGRN* | *# TFs (activators, repressors)* | *# peaks* | *# genes* | *# connections TF-enhancer-genes* |
| --- | --- | --- | --- | --- |
| *Macrophage naive (naive)* | 114 (52, 28) | 1802 | 1793 | 3209 |
| *Macrophage IFN-γ primed (primed)* | 126 (65, 31) | 6742 | 3869 | 22082 |
| *Macrophage infected with Salmonella (infected)* | 92 (35, 26) | 1411 | 1454 | 2128 |
| *Macrophage IFN-γ-primed and infected (primed-infected)* | 78 (30, 22) | 5184 | 2732 | 14697 |
| *AML* | 53 (30, 4) | 2896 | 2525 | 5466 |
| *Primary CD4+ T-cells* | 94 (20, 16) | 3469 | 3258 | 8920 |

As part of GRaNIE, TFs are classified as activators or repressors based on whether the level of TF RNA is positively or negatively correlated with the accessibility surrounding its motifs as originally proposed in (Berest et al. 2019). We observed a slightly larger number of TFs classified as activators than as repressors (1.5-2 fold), yet activators tend to be connected with more peaks resulting in over 10-fold more peaks being linked to an activator than a repressor **(Fig. S5A-D; Table 1).** Notably, in all eGRNs, only about 20-30% of the peaks are linked to their closest gene TSS **(Fig. 1G),** an observation that is consistent with previous observations in pulmonary arterial endothelial cells (7) and iPSC derived cardiomyocytes (31).

On average, a gene is linked to 4.4 (naive), 2.9 (infected) and 5.9 (primed) peaks, of which 1.8 (naive), 1.5 (infected) and 5.7 (primed) are TF-bound and thus part of a GRaNIE eGRN **(Fig. 1H, Fig. S7).** This discrepancy suggests that we are still missing some TF-peak interactions (see limitations). The majority of TFs are connected to very few genes, yet a handful of TFs are connected to over 50 genes, as exemplified for the eGRN from naive and the infected macrophages **(Fig. 1I),** which is in line with the typically scale-free structure of GRNs (32). The most connected TFs differ between the infected and the naive eGRN and include many well-established macrophage TFs such as IRF8, NFkB2, and RELB (33,34) and also less well established TFs such as MBD2, FOSL1 and NRF1, which have been implicated in macrophage biology in more recent studies in mice provide evidence for a role of MBD2, NRF1 FOSL1 in macrophages (35–37).

To dive into the biological processes captured by eGRNs, GRaNIE provides functionalities for identifying subnetworks, or communities (using Louvain clustering by default, as implemented in the *igraph* package in R (38)), and performing GO term enrichment on them. In line with a scale-free architecture of the networks we typically observe a few large communities and a long tail of very small and isolated nodes for each eGRN **(Fig. S8).** Among the communities **(Fig. S8A)** of the naive macrophage eGRN, one is enriched for GO terms related to proinflammatory processes (response to IL-1, chemotaxis, response to IFN-γ) and one for anti-inflammatory processes (angiogenesis, cytoskeleton reorganisation, positive regulation of osteoblast differentiation; **Fig. 1J),** thus recapitulating the potential of naive macrophages to polarise into either M1 (pro-inflammatory) or M2 (anti-inflammatory) cell states (39). We find the M1-phenotype cluster regulated by NFkB1/2 and REL, while the M2-phenotype cluster is regulated by TEAD1/2 and GATA4. Among the communities of the infected macrophage eGRN **(Fig. S8D),** one was enriched for proinflammatory processes, one for phagocytosis related processes, and one for chemotaxis **(Fig. 1K),** thus recapitulating the most important faces of macrophage function (40,41), (42). Notably, each of these functional communities was regulated by a specific set of TFs: IRF8 for the proinflammatory community, MBD2, NFR1, and ETV1 for the phagocytosis, and MECP2for the chemotaxis.

As utility evaluation of the GRaNIE eGRNs, we compared the GO terms that were enriched for each of the TF regulons to terms enriched in “randomised regulons” (same genes randomly grouped into sets of the same size; see Methods). These randomised regulons were enriched in less specific GO terms unrelated to macrophage biology compared to the real regulons **(Fig. S9).**

In summary, these results demonstrate that GRaNIE-inferred eGRNs capture molecular evidence from eQTLs, ChIP-seq and CAGE data, and are useful for investigating TF-driven biological processes in a cell type/state-specific manner.

### Conceptual description of GRaNPA, an approach for evaluating the biological relevance of GRNs and TFs

To evaluate the biological relevance of GRNs and estimate the context-dependent importance of specific TFs, we propose a test based on the premise that cell-type specific GRNs should capture TF-driven gene expression regulation in a cell type specific manner. We devised a machine learning approach, GRaNPA (Gene Regulatory Network Performance Analysis), that evaluates how well the bipartite TF-gene connections of an eGRN can predict cell-type specific differential gene expression and at the same time identifies the TFs that are important for the prediction.

GRaNPA requires differential RNA expression data for the cell type for which the GRN was constructed that is independent from the data used to generate the GRN. It then trains a random forest regression model to predict a differential expression value per gene, based on the TF-target gene connections from the GRN in a 10-fold cross validation setting (see methods; **Fig. 2A).** To assess the specificity of the network, it also trains a separate model based on randomized edges of similar structure and compares the obtained predictions (using R^2^, AUPRC and AUROC). A performance greater than 0.05 (R^2^) for the random networks indicates that even unspecific TF-gene connections can predict differential expression and thus serves as a network specificity-control. Lastly, to assess overfitting, GRaNPA trains the same random network on completely random differential expression data (uniform distribution between −10 to 10; see Methods). A performance greater than 0.05 for a random network to predict random values indicates that the model is overfitting. Notably, GRaNPA can be applied to assess any GRN that contains TF-gene connections and may be used to benchmark GRNs constructed using various methods (see below).

**Figure 2:**
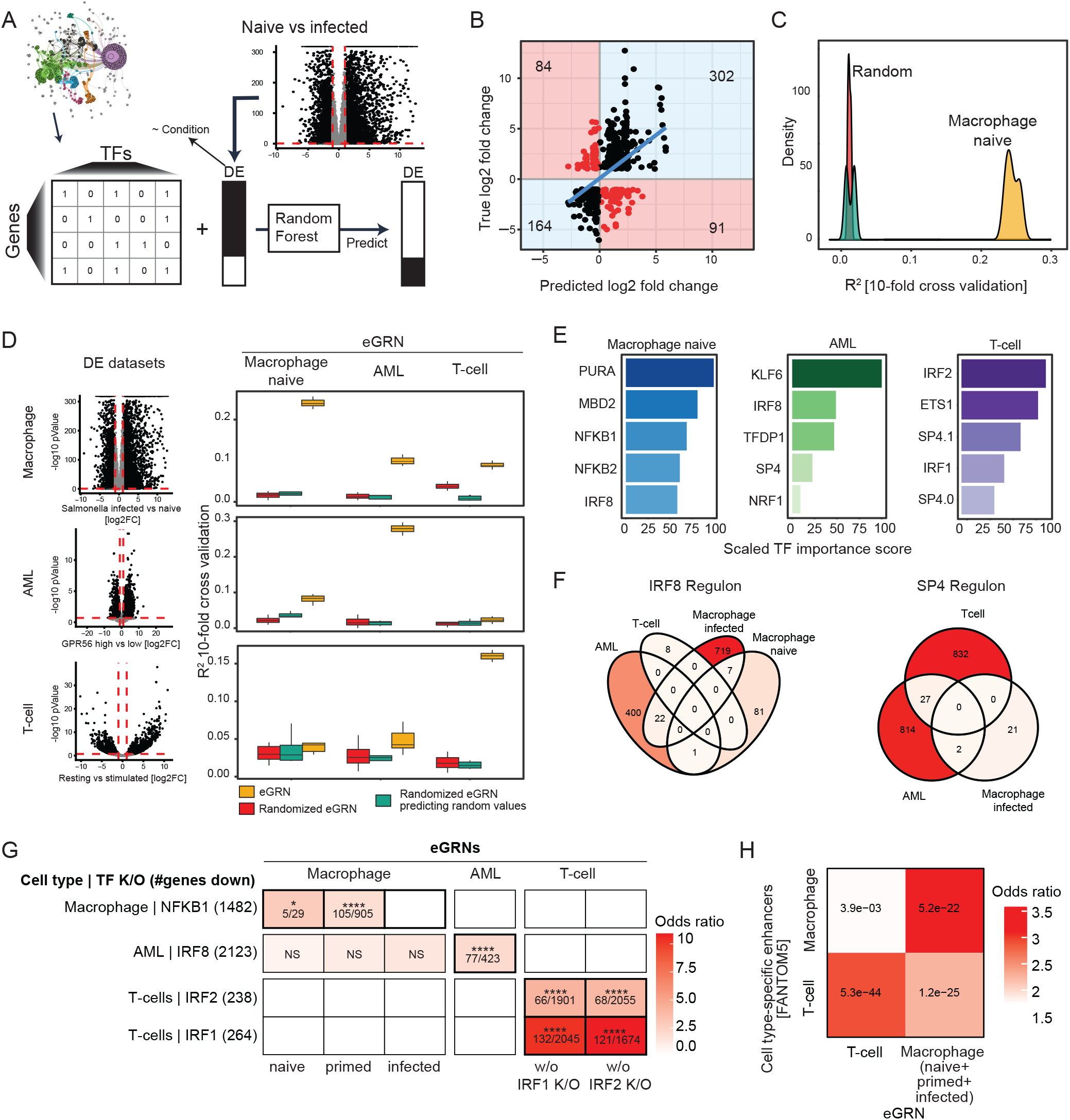
Overview and application of GRaNPA. **A)** Schematic of the general GRN evaluation approach GRaNPA. B) Output of GRaNPA is shown as true vs predicted log2 fold-changes for the macrophage expression response to salmonella infection. Predictions are based on the naive macrophage eGRN (see **Fig. S10** for the other macrophage eGRNs). C) Output of GRaNPA is shown as density distribution of R^2^ for ten random forest runs for the naive macrophage eGRN predicting differential expression upon salmonella infection, along with the two randomised controls. D) GRaNPA evaluation of eGRNs for naive macrophages (left), AML (middle) and T-Cells (right) of differential expression from macrophages infected with Salmonella vs naive (top), two subtypes of AML (middle), and resting vs stimulated T-cells (bottom). Red lines indicate the log2 fold-change (vertical line) and p-value (horizontal line) thresholds for genes included in the GRaNPA analysis. E) Top 5 most important TFs for each of the eGRNs in D) based on prediction in the same cell-type. **F)** Overlap of SP4 (left) and IRF8 (right) regulons between eGRNs from different cell types (only eGRNs with at least one connection to the respective TF are shown). **G)** Enrichment (odds ratio - OR) of NFKB1, IRF8, IRF1 and IRF2 target genes identified in cell-type specific knock/outs (K/O, rows) in the matching macrophage, AML and T-cell eGRN regulons (columns). Numbers in cells indicate: “#genes in regulon and down in TF K/O” / “#genes in regulon”. Asterisks indicate significance of the enrichment (NS: non-significant, *: p-adj. <0.05, ****: <0.0001). H) Enrichment of T-cell and macrophage-specific FANTOM5 CAGE enhancers among the T-cell and macrophage eGRN peaks. The numbers inside the tiles are BH-adjusted p-values. The macrophage eGRN is the union between the infected, naive and primed eGRNs.

In short, our strategy is based on the following steps:

1. Obtain differential expression data for cell types matching the GRNs
2. For each cell type, train a random forest regression model (10-fold cross validation) to predict a differential expression value per gene based on TF-gene links from the eGRN.
3. Compare the performance of models learned on real and random TF-gene links, and TF-gene links from other cell types (cross-validation R^2^).
4. Identify important TFs for the given differential expression response

Furthermore, given a predictive GRN and specific differential expression data, GRaNPA estimates the importance of each TF towards the prediction, which provides candidate driver TFs for a specific expression response. The calculation of TF importance is based on the in-built *importance* function in the R package *ranger* that quantifies importance of features in random forest models based on how much their exclusion affects prediction accuracy. GRaNPA is implemented as a user-friendly R-package (https://git.embl.de/grp-zaugg/GRaNPA) and documentation is available at (https://grp-zaugg.embl-community.io/GRaNPA/).

### GRaNPA evaluation of the macrophage eGRNs

To evaluate the predictive power of the macrophage eGRNs and identify the TFs driving a specific expression response, we obtained RNA-seq data for naive and Salmonella-infected macrophages from (28), and calculated the differential expression using DESeq2 (43) (Methods). For evaluations, we generally excluded samples that were used for the eGRN reconstruction.

GRaNPA evaluation of the four macrophage eGRNs with these data revealed that three of them perform well in predicting the differential expression values (regression) measured as R^2^ (0.15-0.25; **Fig. 2B-C, Fig. S10,11)** and direction of change (classification) as measured by the area under the precision-recall and receiver operating curves (AUPRC: 0.71-0.88; AUROC: 0.65-75; **Fig. S12,13),** respectively. The performance of the corresponding randomised networks was <0.05 R^2^ for two of them **(Fig. 2B-C, Fig. S11-13).** The eGRN for primed-infected macrophages was unable to predict any differential expression **(Fig. S10-13),** which is in line with the failed ChIP-seq validation **(Fig. S2)** and shows the concordance of GRaNPA evaluation with molecular evidence. The significant difference between the randomised and the actual networks shows that the eGRNs indeed capture biologically relevant links between TFs and genes.

### eGRNs built from single cell types show cell-type specific predictions

We next assessed the cell-type specificity GRaNIE-inferred eGRNs. To this end, we obtained datasets in different cell types with matched RNA and chromatin accessibility data for primary human CD4+ T-cells (44) and from acute myeloid leukaemia (AML) (45) and (46). We ran GRaNIE using the same parameters as described above and obtained additional eGRNs for primary CD4+ T-cells (Table S5) and for AML (Table S6).

To assess their cell-type specific prediction power, we ran GRaNPA on the naive macrophage, the T-cell, and the AML eGRNs, and compared their performance to predict differential expression in each of the three cell types. Specifically, we quantified differential expression between resting and lipopolysaccharide (LPS) stimulated follicular CD4+ T-cells (data from (47)), between two subtypes of AML (GPR56-high vs GPR56-low; data from (45)), and between naive vs salmonella-infected macrophages (data from (28)). We found that the eGRN that matches the respective cell type led to the best prediction **(Fig. 2D).** While T-cells and macrophages were only predictive in their own cell type, the AML eGRN was to a smaller extent also predictive for the macrophage response. Since AML cells and macrophages are both from the myeloid lineage this could indicate some shared regulatory architecture between them. Notably, we found that the R^2^ values can be boosted by adding gene specific features, such as the expression variation across individuals, in line with our previous work (48) **(Fig. S14).** As we are here mostly interested in evaluating TF-gene links, GRaNPA doesn’t add gene-specific features by default.

Using the TF-importance estimation implemented in GRaNPA, we observed that among the top 5 important TFs most are unique for one cell type **(Fig. 2E)** with the exceptions of IRF8, which was important in AML and macrophages, and SP4, important for both AML and T-cells. Notably, the IRF8 regulons (defined as all genes connected to a TF) in AML and macrophages had only 22 genes (and no single enhancer) in common, while each included hundreds of non-overlapping genes **(Fig. 2F).** Similarly, the SP4 regulons of T-cells and AML were almost mutually exclusive. This suggests a highly cell-type specific regulon composition of IRF8 and SP4.

As an orthogonal validation of the cell-type specificity of the TF regulons from GRaNIE eGRNs, we compared the regulons with differential expression data upon TF knockout (K/O) in the same cell type. We obtained data for one or two of the top five important TFs in each cell type: NFKB1 in macrophages (49), IRF8 in AML (50) and IRF1 and IRF2 in T-cells (44). The genes downregulated upon TF K/O were significantly enriched in the TF regulons of the respective cell types **(Fig. 2G).** Notably, genes downregulated upon IRF8 K/O in AML were specifically enriched for the IRF8 regulon in AML, despite the fact that IRF8 is also an important TF in macrophages **(Fig. 2E).** This suggests that the cell-type specificity of the GRaNPA predictions is not only dependent on distinct sets of TFs driving the response, but also on the genes the TF regulates in that cell type, highlighting the importance of cell type-specific eGRNs.

To validate the cell-type specificity of enhancers in GRaNIE we obtained cell-type specific enhancer maps from FANTOM5 using CAGE data for T-cells and macrophages (30). Quantifying their overlap with enhancers from T-cell and macrophage eGRNs revealed a stronger significant enrichment among the enhancers from the same cell type as compared to opposite cell types **(Fig. 2H).**

The eGRNs connect TFs to genes through active regulatory regions, comprising both promoters and enhancers. The predictive evaluation setup allowed us to compare the relative importance of promoter (i.e. <10kb from TSS) and enhancer links (>10kb from TSS) in different eGRNs. To do so, we divided the gene-peak pairs into ten groups based on their distance to the TSS and ran GRaNPA for each group separately. The promoter-only eGRNs from infected and primed macrophages showed limited or no predictive power **(Fig. S15).** This highlights the importance of enhancers and is in line with a recent study that demonstrated the importance of considering enhancers for predicting the cell-fate potential of TFs (5).

### Application of GRaNPA to compare GRaNIE eGRNs with other GRN methods

Notably, GRaNPA is applicable to assess any type of bipartite TF-gene network and can be used more generally to assess the utility of a GRN for understanding cellular response to specific perturbations. Here we used it to evaluate the performance of several previously published TF-gene GRNs that draw links between TFs and genes based on different approaches: DoRothEA, which uses manual curation combined with a data-driven approach including co-expression to draw TF-gene links (15,51,52), ChEA3, which uses ChIP-seq experiments from ENCODE, ReMap, or literature to draw TF-gene links (17), RegNet, a curated network integrating TFs and miRNAs (18), and TRRUST, which is a curation of TF-gene links based on PubMed indexed articles (16). We also included an enhancer-based GRN inferred with ANANSE from macrophage data.

The cell-type matched GRaNIE eGRNs and DoRothEA ABC showed good prediction for all datasets tested. The TRRUST, RegNet and ChEA3 networks showed slight predictive power for macrophages, while the only other enhancer-based network ANANSE showed very poor performance across all cell types **(Fig. 3A).** Thus, GRaNIE networks outperformed most other networks and was on par with the highly curated DoRothEA.

**Figure 3:**
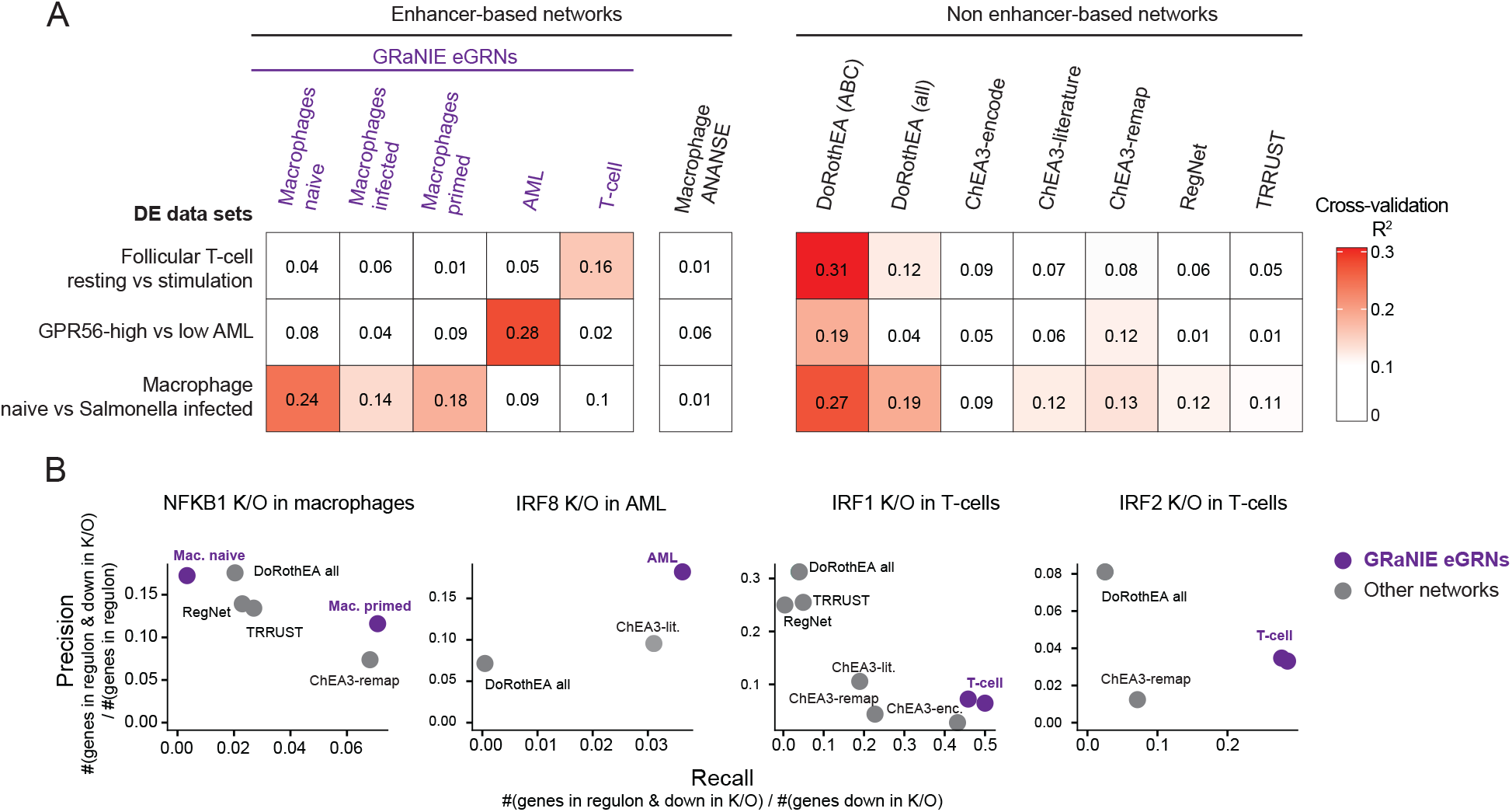
Evaluation of GRaNIE eGRNs and other GRN approaches. **A)** GRaNPA evaluation of five GRaNIE eGRNs (macrophage naive / primed / infected, AML, and T-cells), another enhancer-based eGRN inferred with ANANSE (5), and publicly available TF-gene networks based on data curation (DoRothEA ABC and all (51)), ChIP-seq data (ChEA3 encode, literature, and ReMap (17)), manual curation (TRRUST (16) and REGNET (18)). GRNs are evaluated by GRaNPA for their performance in predicting the differential expression of resting vs stimulated follicular T-cells, GPR56 high vs low AML, and naive vs Salmonella-infected macrophages. Numbers in squares indicate R^2^ values. **B)** Precision-Recall evaluation of the NFKB1, IRF8, IRF1 and IRF2 regulon from the networks in A) for identifying genes down-regulated upon K/O of the respective TF. For GRaNIE eGRNs (purple), the performance of cell-type matching networks is shown, other networks are the same across all analyses.

To further compare cell-type specificity, we assessed the overlap between the TF-regulons identified in the networks with reasonable predictive power and the genes downregulated upon K/O of the same TF (data introduced in **Fig. 2G).** Overall, the cell-type matched GRaNIE eGRNs outperformed all other networks in terms of recall **(Fig. 3B).** Of note, the absolute recall was rather small, likely owing to the fact that TF K/O has a lot of indirect downstream effects that are not captured by the direct mechanistic links of eGRNs. GRaNIE also outperformed all other networks in terms of precision in AML. While DoRoThEA achieved the highest precision for IRF1 and IRF2 K/O in T-cells, the recall was smaller. Overall, this cell-type specific TF K/O evaluation highlights the importance of unbiased and cell type-specific eGRNs.

### Macrophage eGRNs reveal distinct set of TFs driving response to different types of infection

We next highlight the biological insights that GRaNIE and GRaNPA can provide. Specifically, we employed them for studying different types of pro-inflammatory (M1) response of macrophages to bacterial infections as well as a more M2-like response of breast cancer associated macrophages. We obtained data from previously published studies that measured the expression response of macrophages infected with *Mycobacterium Tuberculosis* (MTB) (53), *Listeria monocytogenes* (54), *Salmonella Typhimurium* (28,54)), and stimulation with IFN-γ (28), and a study that compared tumour associated macrophages with tissue-resident macrophages from breast cancer tissue (55). The studies are quite diverse in their set-up **(Table 2).**

**Table 2:** Differential expression experiments for the different infection settings.

Differential expression summary for the different infection datasets / cell types, their treatments and the comparisons used for the differential expression analyses.
| Cell types | Treatment | Comparison | Reference |
| --- | --- | --- | --- |
| Monocyte-derived macrophages from healthy donors | Listeria monocytogenes | Uninfected vs 2 hours post infection | (54) |
|  | Salmonella Typhimurium |  |  |
|  | Mycobacterium Tuberculosis strain resistant to BDQ treatment | Uninfected vs 18h post infection | (53) |
|  |  | Uninfected vs 36h post infection |  |
| iPSC-derived macrophages | Salmonella typhimurium | Uninfected vs 5h post infection | (28) |
| | | 18h IFN-γ-primed vs 18h IFN-γ-primed + 5h post infection | |
| | Interferon-gamma stimulation | naive vs 18h IFN-γ-g treatment | |
| Tumour-associated and tissue resident macrophages from human breast tissue | Tumor vs tissue-resident | Tumour vs respective tissue resident macrophages | (55) |

To understand how macrophages respond to these distinct perturbations, we employed GRaNPA using the union of the naive and infected eGRNs **(Fig. S16; Table S4),** which showed good predictions for all conditions **(Fig. 4A),** and determined the important TFs for each response. One of the well-understood responses of macrophages is the IFN-γ mediated activation of the NFKB family of inducible transcription factors (56). In line with this, we find NFkB2 as one of the most important TFs upon IFN-γ stimulation **(Fig. 4B).** The NFkB2 regulon was enriched for GO terms related to chemokine signalling and taxis **(Fig. S17)** and strongly upregulated in response to IFN-γ **(Fig. 4C),** demonstrating the ability of GRaNPA to identify biologically meaningful TFs. To assess the robustness of GRaNPA we compared the TF importance predictions across two independent datasets from Salmonella infected macrophages, which revealed very similar profiles despite differences in the experimental set-up **(Fig. 4B;** iPSC-derived vs monocyte-derived macrophages) and time points (5h and 2h post infection), thus highlighting the robustness of GRaNPA as well as the biological congruence between the experiments.

**Figure 4:**
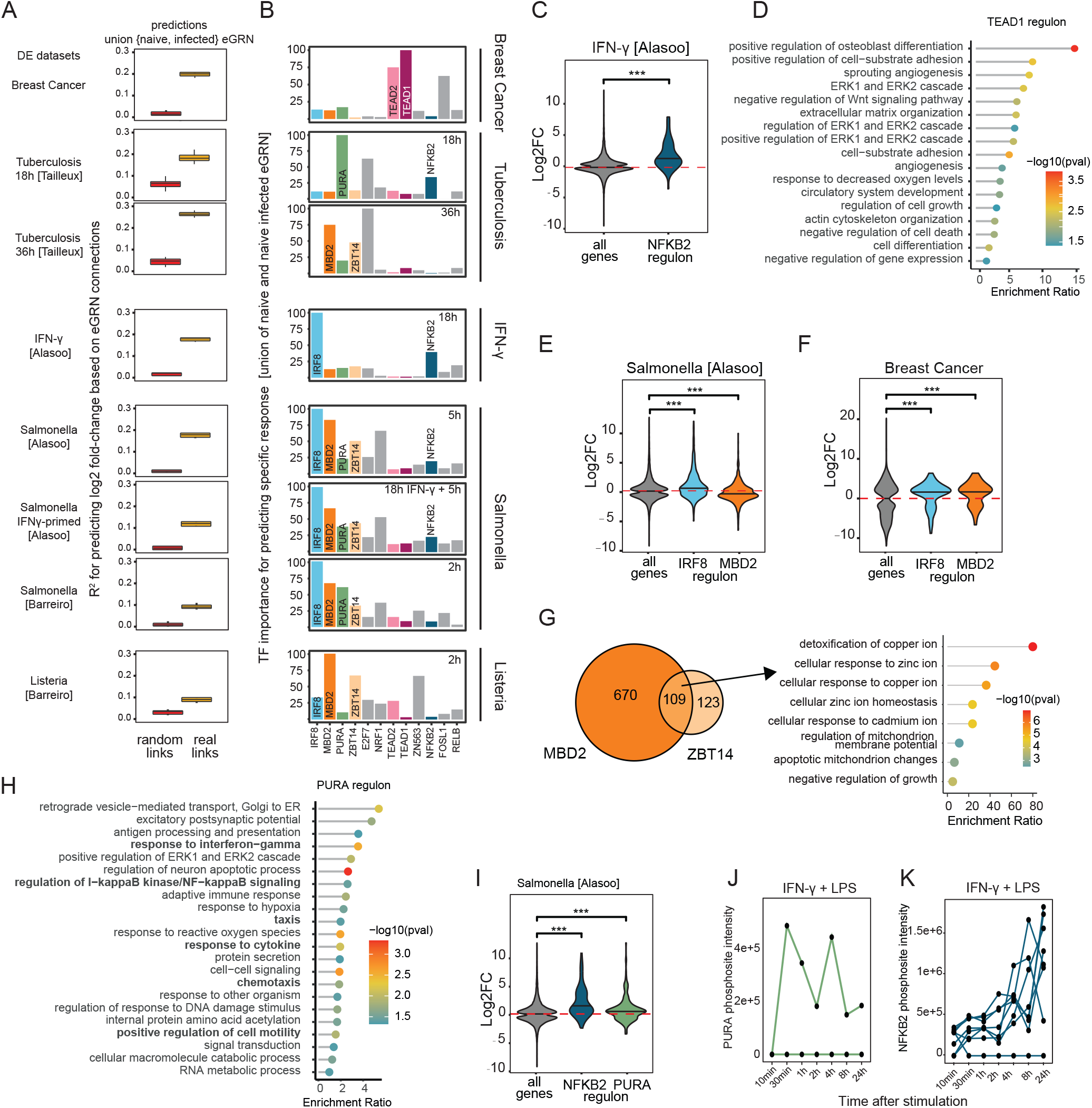
Application of GRaNIE and GRaNPA to investigate macrophage biology. **A)** GRaNPA evaluation of the union of the naive and infected macrophage eGRNs (naive+infected eGRN; real links) and the corresponding randomized control network (random links) across eight experimental settings of macrophage perturbations. B) TF importance profiles for each of the eight infection settings from A. The top 5 most predictive TFs in any of the settings are displayed. TFs discussed in the text are individually labelled and colored. C) Distribution of log2 fold-changes for genes in the NFKB2 regulons from the naive+infected eGRN (dark blue) are shown for IFN-γ stimulation vs naive macrophages alongside the response of all genes (grey). D) GO enrichment of the TEAD1 regulon. E-F) Distribution of log2 fold-changes of Salmonella infection vs naive macrophages (E) and for breast-cancer associated macrophages (F) are shown for genes in the IRF8 (blue) and MBD2 (orange) regulons alongside the response of all genes (grey). G) The overlap between the MBD2 and ZBT14 regulons are shown as Venn Diagram (left). Enriched GO terms for the genes in the intersection are shown as lolli plot (right). H) GO enrichment of the PURA regulon. I) Distribution of log2 fold-changes of Salmonella infection vs naive macrophages for genes in the NFKb2 (blue) and PURA (green) regulon alongside the response of all genes (grey). J-K Normalised mass spectrometry intensity values (y-axis) for phosphosites detected on PURA (green, J) and NFKB2 (blue, K) in macrophages cultured in the presence of M1 polarising stimuli (IFN-γ and LPS) for indicated time points (x-axis). Lines show individual phosphosites detected on each respective TF.

In contrast, TF-importance profiles were highly variable across conditions **(Fig. 4B),** likely reflecting different roles of macrophages (M1 vs M2) and their variable defence mechanisms triggered by the pathogens (57). Breast-cancer associated macrophages showed the most distinct profile with TEAD1/TEAD2 as important TFs. GO analysis of the TEAD1/2 regulon revealed a strong enrichment for angiogenesis, osteoblast differentiation and ERK signalling among others **(Fig. 4D),** in line with a more M2-like phenotype (58,59)

The most important TF for predicting the response to Salmonella infection was IRF8, followed by MBD2 and ZBT14 **(Fig. 4B).** IRF8 is a known pro-inflammatory interferon response factor, associated with the pro-inflammatory (M1) polarisation of macrophages (60), which we confirmed in our data using gene-set enrichment analysis of the IRF8 regulon **(Fig. S18).** Less is known about MBD2 and ZBT14 in macrophages, although MBD2 has been linked to intestinal inflammation in mice (35) and with an M2 macrophage programme in pulmonary fibrosis (61). In line with this, the MBD2 regulon was down-regulated in response to infection **(Fig. 4E, S13)** but upregulated in breast cancer associated macrophages **(Fig. 4F),** showing the opposite pattern to the IRF8 regulon **(Fig. 4E-F).** We further find an enrichment of the M2 gene set among the MBD2 regulon in breast-cancer associated macrophages **(Fig. S18).** The MBD2 and ZBT14 regulons show significant overlap **(Fig. 4F,** p=3.3e-13, hypergeometric test) and genes jointly regulated by them are enriched for terms related to response to metal ions **(Fig. 4G).** The use of zinc and copper ions in macrophage defence strategies is well documented (62,63) and given that the ZBT14 and MBD2 are only important for predicting response to pathogens, and not for predicting response to IFN-γ stimulation **(Fig. 4B),** we speculate that ZBT14 and MBD2 may jointly induce a macrophage-intrinsic mechanism to counteract toxic metal ions, potentially aimed at overcoming the toxic effects of its own weapons.

### GRaNPA identifies PURA as putative proinflammatory TF in macrophages

Among the TFs that are less well known for their role in macrophages we find PURA for many of the infection settings. In line with a pro-inflammatory role of PURA we found GO terms associated with chemotaxis and IFN-g response enriched among genes in its regulon **(Fig. 4H).** Gene set enrichment analysis (GSEA) found the M1 gene set significantly enriched among the PURA-regulated differentially expressed genes in Salmonella **(Fig. S18).** Furthermore, the expression of genes in the PURA regulon followed the same pattern as the regulon of the pro-inflammatory NFkB2: upregulated upon Salmonella infection **(Fig. 4I).**

To follow-up on a potential role of PURA in M1 polarisation we obtained phosphoproteomics data that was collected upon stimulating macrophages with LPS and IFN-γ towards the M1 phenotype (64). This revealed a specific increase in phosphorylation of Thr187 upon LPS IFN-γ **(Fig. 4J),** following a similar pattern as the phosphorylation pattern of NFkB2 **(Fig. 4K).** Notably, the infection-responsive phosphosite in PURA is located in the Pura repeats region, which is implicated in DNA binding and crucial for PURA function (65). Phosphorylation of DNA-binding regions has been associated with activation of other TFs (66), suggesting that activation of PURA is perhaps important for M1 polarisation, providing further evidence for its role in macrophages’ pro-inflammatory response.

Overall, these results highlight the use of GRaNPA in conjunction with cell-type specific eGRNs for investigating the biological functions that are regulated by a TF in a specific cell type.

### Macrophage-specific eGRNs are enriched in fine-mapped GWAS variants and immune-related traits

The strength of the eGRN framework is that we can specifically investigate the role of gene regulatory elements such as enhancers, which are enriched for disease-associated genetic variants (1). We therefore sought to explore the macrophage eGRNs to learn about gene regulatory mechanisms underlying associations of genetic variants with common complex traits and diseases.

First, we tested whether the peak regions specific to the three macrophage eGRNs that GRaNPA identified as predictive in at least one infection setting (naive, primed, infected) were enriched in heritability for 442 GWAS traits **(Table S9).** We applied *stratified linkage disequilibrium score regression* (S-LDSC (67)) and compared the eGRN peaks to all peaks identified in macrophages (see Methods). Notable enrichments include HbA1c measurement, a measure for diabetes severity, large artery stroke and heart failure for the naive eGRN; adolescent idiopathic scoliosis and nonischemic cardiomyopathy for the primed eGRN and rheumatoid arthritis (RA) and systemic lupus erythematosus (SLE) for the infected eGRN **(Fig. 5A).** SLE and RA are both driven by activated macrophages (68,69) as a result of known (for SLE) or hypothesised (RA) upregulation of IFN-γ signalling (70–72). Interestingly, we find enriched heritability for these traits in the peaks for the IFN-γ primed eGRN, but not for either naive or infected eGRNs. Given this association, we also assessed the heritability enrichment of the regulatory elements and genes connected to the TFs that are particularly important for predicting the response of macrophages to IFN-γ (NFkB1/2, RELB, IRF8). Inflammatory bowel disease (specifically ulcerative colitis) comes out as the top enriched trait **(Fig. S19),** which is in line with the known role of IFN-γ in this disease (73). Literature evidence for other traits is summarised in **Table S10.**

**Figure 5:**
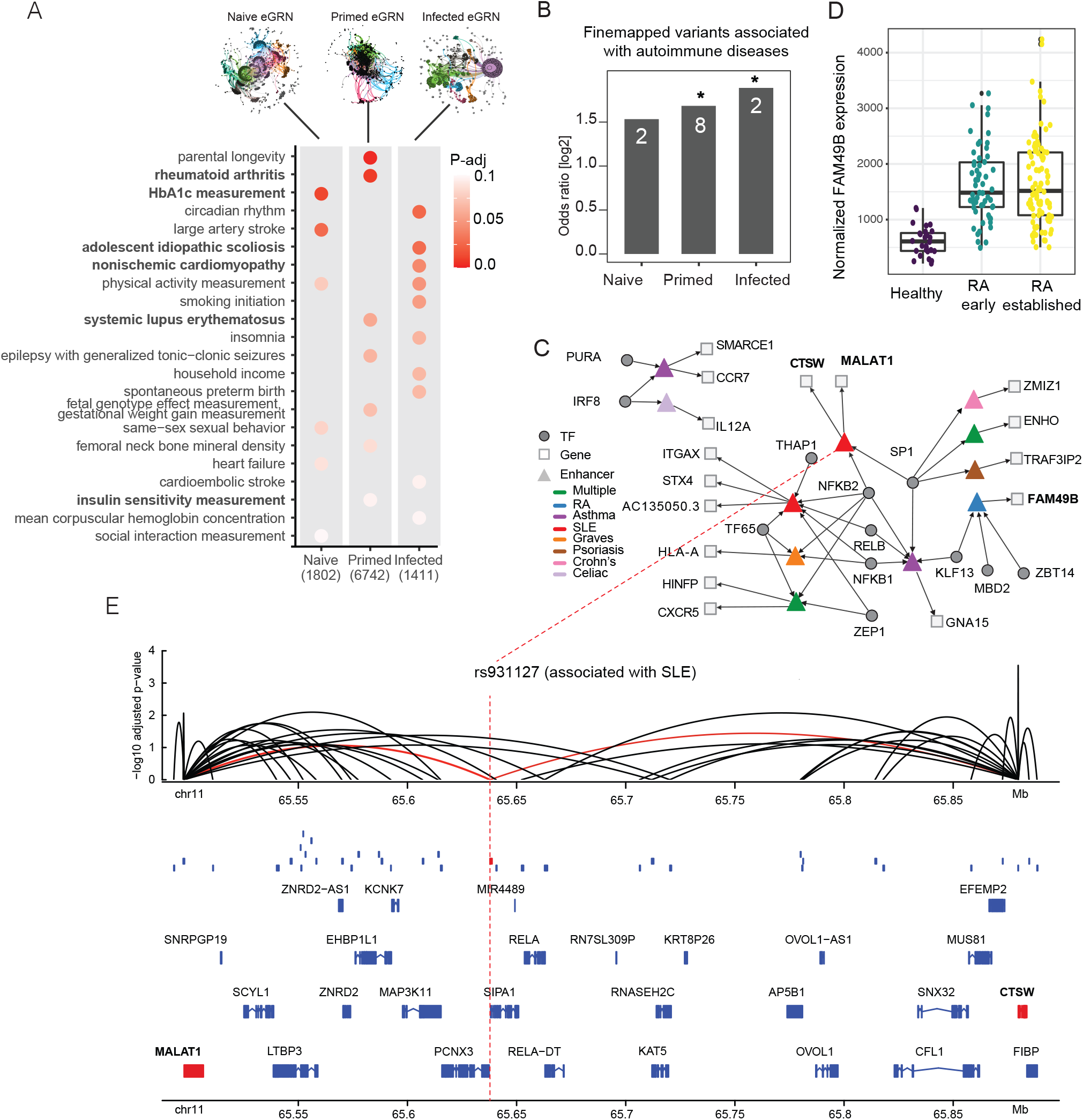
Application of GRaNIE for investigating trait-associated SNPs. **A)** Heritability enrichment is shown for the naive, primed and infected macrophage eGRNs. The p-value is adjusted within each trait. B) The enrichment of fine-mapped GWAS variants within the naive, primed, and infected eGRNs is shown as odds ratios; *: p-value <0.05; n: number of SNPs. C) The tripartite TF-enhancer-gene network involving all fine-mapped GWAS variants for autoimmune diseases. D) Normalized expression level of FAM49B is shown as boxplot for healthy controls and from synovial tissue of patients suffering from early (green) and established (yellow) rheumatoid arthritis (RA). Data from (74). E) The genomic context of the fine-mapped, SLE-associated variant rs931127 in a ATAC-seq peak (red box) as gene tracks, including other peaks present in the infected macrophage eGRN (blue boxes), and peak-gene links from the infected macrophage eGRN (arcs). Genes targeted by the peak overlapping with rs931127 (red) are colored in red.

Next, we zoomed in to a specific set of fine-mapped GWAS variants associated with autoimmune diseases with a known link to macrophages. Across all three macrophage eGRNs we found in total 11 unique fine-mapped variants that were located in the regulatory regions (2 in the naive, 2 in infected, and 8 in the primed eGRN). The infected and primed eGRNs were significantly enriched for containing fine-mapped variants **(Fig. 5B).** Investigating the TFs regulating the fine-mapped auto-immune disease enhancers, we find the known immune response TFs NFKB1/2, RELB and IRF8, but also MBD2, ZBT14 and PURA **(Fig. 5C),** which we identified as important TFs for predicting response to infection. One gene linked to a fine-mapped SNP that is regulated by MBD2 and ZBT14 is FAM49B. Inspection of FAM49B expression in synovial tissue in a cohort of RA patients compared to healthy controls (74) revealed a misregulation upon disease onset, providing additional evidence that FAM49B is indeed the gene targeted by the fine-mapped SNP **(Fig. 5D).**

One of the fine-mapped SNPs in SLE is rs9893132, located in an enhancer regulated by SP1 and NFKb2 that is linked to the non-coding RNA MALAT1 and the gene cathepsin W (CTSW) in the primed macrophage eGRN. Both genes are over 50kb from the fine-mapped SNP and there are several other genes in the locus that are not linked to the enhancer in question **(Fig. 5E).** MALAT1 has been implicated in SLE through several studies (75), suggesting that the fine-mapped SNP may target MALAT1. While CTSW expression in lymphocytes has been linked to autoimmune diseases (e.g. (76)), its role in macrophages is much less studied. Yet CTSW knockdown in macrophages reportedly increased *Mycobacterium tuberculosis* survival in macrophages (77), suggesting it does play an important function in the pro-inflammatory macrophage response. Overall, the macrophage eGRNs provide the putative target genes of 11 fine-mapped GWAS loci, often linking to genes that are over 50kb away from the SNP **(Table 3; Fig. S20).**

**Table 3:** Predicted target genes of fine-mapped GWAS autoimmune variants using macrophage eGRNs.

Fine-mapped GWAS variants for autoimmune diseases generated using probabilistic identification of causal SNPs (PICS; see Methods) for *hg38* build overlapping with the peaks of macrophage eGRNs.
| Peak | TF | Gene name | FM.gwas | rsid | Disease | Network |
| --- | --- | --- | --- | --- | --- | --- |
| chr8:129939305-129940672 | KLF13.0.D, MBD2.0.B, ZBT14.0.C | FAM49B | chr8:129939865 | rs11785995 | Rheumatoid arthritis | naive |
| chr17:40598595-40599644 | PURA.0.D | SMARCE1 | chr17:40598769 | rs9893132 | Asthma | naive |
| chr19:3135922-3136231 | KLF13.0.D, NFKB1.1.B, NFKB2.0.B, RELB.0.C, SP1.0.A, SP1.1.A | GNA15 | chr19:3136093 | rs117552144 | Asthma | primed |
| chr16:31265059-31265802 | NFKB1.1.B, NFKB2.0.B, RELB.0.C, TF65.0.A, THAP1.0.C, ZEP1.0.D | STX4, AC135050.3, ITGAX | chr16:31265490 | rs1143679 | Systemic lupus erythematosus | primed |
| chr6:30006131-30007036 | NFKB1.1.B, NFKB2.0.B, TF65.0.A | HLA-A | chr6:30006148 | rs4313034 | Graves' disease | primed |
| chr11:65637802-65638765 | NFKB2.0.B, SP1.0.A, SP1.1.A | MALAT1, CTSW | chr11:65637829 | rs931127 | Systemic lupus erythematosus | primed |
| chr11:118883323-118883647 | NFKB2.0.B, TF65.0.A, ZEP1.0.D | CXCR5, HINFP | chr11:118883644 | rs630923 | Multiple sclerosis, Inflammatory bowel disease, Crohn's disease | primed |
| chr6:111605185-111606373 | SP1.0.A | TRAF3IP2 | chr6:111605706 | rs7769061 | Psoriasis | primed |
| chr9:34709959-34710335 | SP1.0.A, SP1.1.A | ENHO | chr9:34710263 | rs2812378 | Rheumatoid arthritis, Celiac disease | primed |
| chr10:79285352-79285717 | SP1.1.A | ZMIZ1 | chr10:79285450 | rs1250569 | Crohn's disease | primed |
| chr10:79285352-79285717 | SP1.1.A | ZMIZ1 | chr10:79285523 | rs1250568 | Celiac disease | primed |
| chr17:40598595-40599644 | IRF8.0.B | CCR7 | chr17:40598769 | rs9893132 | Asthma | infected |
| chr3:159929439-159930124 | IRF8.0.B | IL12A | chr3:159929885 | rs17753641 | Celiac disease | infected |

## Discussion

Phenotypic variation across individuals has two major sources. On the one hand it is genetic variation that can lead to variation in molecular phenotypes that impact on complex traits. On the other hand, phenotypic variation can arise from variation in the environment that can be long-lived (epigenetics) or short-lived (signalling). Thus, to understand mechanisms underlying phenotypic variation (including disease phenotypes), it is crucial to study the interplay between genetic variants, epigenetic marks and extrinsic cellular signalling. Here we present GRaNIE and GRaNPA, a tool-suite that provides a framework for jointly analysing these three layers and investigating their biological relevance.

GRaNIE is a flexible and user-friendly R/Bioconductor package for building enhancer-based GRNs, while GRaNPA is a separate and independent R package for evaluating the biological relevance (i.e. predictive power) of any general TF-gene networks. GRaNIE requires RNA-Seq and open chromatin data such as ATAC-Seq or ChIP-Seq for histone modifications (e.g, H3K27ac) across a range of samples (mostly tested in a cohort of at least 10-15 individuals), along with TFBS data (that can either be obtained from the package or provided by the user) to generate cell-type specific eGRNs. It provides a range of quality control plots and functionalities for downstream analyses such as identification of communities within the network, and GO enrichment analyses. A dedicated website accompanies the package and is automatically updated whenever a new package version becomes available.

GRaNPA requires a bipartite TF-gene network and gene-specific differential expression values as input, and assesses its predictive power. As a downstream functionality it provides quantification of TF importance for driving a specific differential expression response. Notably, the purpose of GRaNPA predictions is to assess the TF-gene connectivities, and while including additional gene-specific features may improve prediction performance, as shown in (48), it would not help in the assessment of GRNs. An additional advantage of GRaNPA is we can directly compare different GRNs and get a quantitative assessment of their performance for a perturbation of interest, without the need for a “ground-truth” network to evaluate their edges and connectivities.

Compared to most of the available GRNs reconstruction approaches, GRaNIE infers enhancer-based regulatory networks that only captures TF-gene links mediated by enhancers. This has several advantages: first, we showed that at least for macrophages, parts of their expression response to infection could only be predicted when using enhancer connections. Second, including enhancers in TF-driven GRNs allows the investigation of mechanisms underlying GWAS traits that are driven by specific TFs, and facilitates interpretation of (fine-mapped) trait-associated SNPs. Third, since enhancers tend to be highly cell-type specific (78), eGRNs are likely more cell type specific than TF-gene networks. Finally, by requiring a correlation between the expression level of a TF and the accessibility of the peak, in addition to the motif presence, the GRaNIE approach circumvents the inherent limitation of TF binding site predictions, which cannot distinguish between TFs with similar binding motifs (79). This will exclude many TF-enhancer links that have the TF motif yet show no correlation with the TF expression level and are thus likely false positive binding sites in the given cell type. GRaNIE bears conceptual similarity with a method published earlier (22), however the data of this work are not available anymore and the software is neither maintained nor cannot be downloaded/used.

eGRNs consist of TFs and their respective downstream enhancer/promoter and gene targets, which means we can zoom in to network communities that capture specific pathways or functions. For example, we showed that when we divide the network into communities, each community is enriched in distinct TF-driven biological processes. The modularity of the eGRN also showed that NFKB1, NFKB2, RELB and IRF8, the TFs important for predicting the macrophage response to IFN-γ priming, and their connected regulatory elements and genes were specifically enriched for heritability of GWAS traits that are commonly linked to IFN-γ signalling. In contrast to other approaches for interpreting trait-associated variants that are solely based on epigenetics such as the activity by contact model (80), or purely based on genetics, such as eQTLs (4) or variable chromatin domains (81), eGRNs by GRaNIE capture TF-peak-gene links based on variation due to genetic, epigenetic, or TF-activity differences across individuals, thus integrating these three layers in one framework. In sum, eGRNs can be used to identify the target genes of individual TFs (regulons), to pinpoint the cell-type specific regulatory regions that connect to a TF, and to investigate genetic variation in the tripartite TF-regulatory element-gene graphs.

Comparing eGRNs across cell types revealed that for some TFs (e.g. IRF8) the regulons are highly cell-type specific. One explanation for this cell-type specific regulons is that TFs may have different co-binding TF partners depending on the cell type. Indeed, in our previous work we found that TFs regulate distinct biological processes depending on their co-binding partners (82,83). An alternative explanation for cell-type specific regulons is that different cell types may differ in their chromatin potential (84).

Among the notable observations from applying GRaNIE and GRaNPA to study the gene expression response in macrophages was that some TFs, including MBD2, were specifically important only for predicting response to bacterial infection, and not for IFN-γ stimulation. Since IRF transcriptional programs are generally more related to a virus response, MBD2 is perhaps needed in the bacterial specific response, indicating that we can potentially use these networks to identify TFs important for responses to different types of pathogens. The observation that GRaNPA identified distinct sets of important TFs for the different responses may reflect that macrophages use several strategies to fight infections, including phagocytosis followed by degradation mechanisms, starvation of pathogens, and recruiting other players in the immune system (57). Another observation is that three of important TFs for predicting the response to infection and but not to IFN-γ are known to bind methylated DNA: MBD2 (85), MECP2 (86), NRF1 (87). Recent reports provide evidence for a pathogen-induced global DNA methylation alteration (88) downstream of NFkB-signalling, and it was shown that MBD2 inhibits IFN-γ by selectively binding to methylated regions in the Statl promoter in other cell types (89). Our results are consistent with a pathogen-response mechanism that is partially mediated by DNA methylation, which may modulate the impact of DNA-methylation sensitive TFs and demonstrates the level of novel biological insights that can be gained with GRaNIE and GRaNPA.

### Limitations of GRaNIE

1. As with all network inference tools, it is important to keep in mind what an edge means. In the case of GRaNIE, the TF-peak and peak-gene links are based on co-variation across biological samples (in this study variation across individuals). Therefore, it will miss links when either of the nodes (TF expression, peak accessibility, or gene expression) is not variable across samples. For instance, if samples are individuals, GRaNIE may miss house-keeping and dosage-sensitive genes, TFs, and enhancers.
2. If GRaNIE is run with ATAC-seq data, the limitations of ATAC-seq apply: i.e. accessibility may not always reflect activity. Generally with ATAC-seq data promoter accessibility is often not strongly correlated with gene expression when considering cells in which the gene is generally active. Therefore, GRaNIE will likely miss some promoter-gene connections. Furthermore, it will not detect TFs that do not affect accessibility.
3. As with most TF-inference based tools, GRaNIE relies on the availability of a TF binding site within a peak. Therefore it will miss TF, for which binding sites are unknown, and TFs binding events that do not rely on the TF motif (e.g. cooperative binding).
4. TF expression is not always predictive of a TF’s role in transcriptional regulation. To circumvent this, GRaNIE offers the option of using TF motif accessibility as an estimate of TF activity. This in turn has the caveat that connections will be based on TF motifs (which can be very similar across TFs).
5. Since GRaNIE is association-based it cannot per se distinguish direct from indirect effects. This is particularly important when running it on samples that are highly variable (e.g. different cell types). It may then become difficult to assess whether the variation in peak accessibility is driven by the TF for which it has a motif, or by some other mechanism. We refer the users to the QC implemented in GRaNIE to judge the extent of such an existing batch effect.
6. The R^2^ values from GRaNPA are often low, even if they are better than those for random networks, suggesting that the eGRNs are not picking up all the signal in the data. Adding gene-specific features e.g. from (48) may substantially improve them if desired.

## Acknowledgements

We thank Anna Mathioudaki for proofreading and for naming GRaNIE and GRaNPA, Martina Muckenthaler, Oriana Marques and Christina Mertens for discussions about macrophage biology. M.K. is funded by the European Union’s H2020 MSCA-ITN Project ENHPATHY, grant agreement number 860002. A.C. is funded by the European Union’s Horizon 2020 MSCA, grant agreement number 847543. ARP has been granted by the Regional Programme of Research and Technological Innovation for Young Doctors UCM-CAM (PR65/19-22460).

## Declarations

We declare that none of the authors have any competing interests.

## Methods

### Datasets used in this study

For all datasets, we performed PCA along with metadata inspection in the PCA space to evaluate whether samples should be discarded as outliers. If we did, we give details in the respective paragraph.

#### Expression and chromatin accessibility data for iPSC-derived macrophages

We used a publicly available dataset (ERP020977) for naive and primed macrophages (iPSC-derived) in two conditions, uninfected and 5-h infected with Salmonella from (28). In total we obtained 304 RNA-seq profiles from 86 different individuals, of which 145 also had ATACseq data available (https://zenodo.org/record/1188300#.X370PXUzaSN). The samples are split into four groups: primed, primed-infected, naive and naive-infected for which 41 (43), 31 (55), 42 (42), 31 (55) paired RNA/ATAC (only RNA-seq) samples were available respectively. The data also contained metadata and peak coordinates. The paired samples were used to reconstruct the eGRNs with GRaNIE (see below). The unpaired RNA-seq data were used for evaluation of the eGRNs with GRaNPA (see below).

#### *Expression data for macrophages infected with Listeria & Salmonella* (GEO accession number: GSE73502)

Pai et al generated expression data on cultured monocytes obtained from PBMCs of healthy donors, for which we downloaded the raw counts data (54). Matured macrophages were divided into three groups: (1) controls and infected by the (2) Listeria and (3) Salmonella bacteria, respectively. We used the RNA-seq data collected 2 hours after infection with Listeria and Salmonella respectively for each of the 57 samples.

#### *Expression data for macrophages infected with Tuberculosis* (GEO accession numbers: GSE133145, GSE143731)

Giraud-Gatineau et al collected two datasets on the effect of bedaquiline (GSE133145) and five other drugs (GSE143731) treatment for *Mycobacterium tuberculosis* infection in Monocyte-derived macrophages from healthy donors (53). The GSE133145 series consists of 16 control and 16 *M. tuberculosis-infected* samples, which are later divided into four groups: untreated / DMSO treatment (control) / two variants of bedaquiline treatment (0.5 or 5 microg/mL). Differential expression analysis revealed that differences caused by treatment are not substantial, so we considered the treatment as a controlling variable. The GSE143731 series consists of 28 control and 28 *M. tuberculosis-infected* samples, which are later divided into groups corresponding to the treatment with isoniazid (INH), rifampicin (RIF), ethambutol, pyrazinamide (PZA) or amikacin (AMK), and control group. We considered treatment as a controlling variable for the differential expression analysis.

#### Expression and chromatin accessibility data for CD4+ T-cells

Paired RNA- and ATAC-seq data were obtained from (44). For RNA-seq, processed count files were obtained from GSE171737. For ATAC-seq, raw sequencing files were obtained from GSE171737 and processed and quality-controlled with an in-house Snakemake pipeline as previously described (27).

#### *Expression data for resting vs LPS-stimulated CD4+ T-cells* (GSE118165)

RNA-seq was obtained from (47), which measured expression in resting and stimulated subsets of CD4-positive T-cells. We used only the naive T-cells for differential expression analyses.

#### Expression and chromatin accessibility for AML

We obtained raw RNA-seq data for 23 AML patients from (90). Processed and quality-controlled ATAC-seq data and peaks for the same patients was obtained from (46).

#### Expression data for TF K/Os

We obtained cell-type specific knock-out (K/O) data for THP1 derived macrophages (NFKB1) (49), the human AML cell line MV4-11 (IRF8) (50), and processed differential expression data from primary human CD4+ T cells (IRF1 and IRF2) (44). For the NKB1 and IRF8 datasets, raw sequencing files were obtained from GSE162015 and GSE163275 respectively, and data processing and quality control was performed with an in-house Snakemake pipeline as described previously (27).

#### Macrophage phosphoproteomics data

Processed quantitative phosphoproteomics data from polarising THP1-derived macrophages was obtained from (46).

### Differential expression analysis (see also Table 2)

Differential expression analysis was performed with DESeq2 (43) for all datasets, typically using the contrast between treatment and no treatment or disease and control. The design formula generally used was therefore “~condition”, unless otherwise stated. Dataset-specific details of the differential expression analysis datasets are described below. As input for GRaNPA, we generally used shrunken log2 fold-changes as implemented in *IfcShrink* from DESeq2 with the *apeglm* method (91) unless otherwise indicated, even though is not a strict requirement of GRaNPA to use any particular transformation.

#### iPSC-derived macrophages infected with Salmonella from (28)

Differential expression was calculated using only the RNA-seq data that was not used for eGRN reconstruction. We quantified differential expression for the following contrasts: naive vs infected, naive vs IFN--γ primed, IFN-γ primed vs IFN-γ primed-infected. The formula used in DESeq2 was “-condition”.

#### Macrophages infected with Listeria and Salmonella from (54)

We analysed the differential expression between control samples, listeria-infected samples and salmonella-infected samples separately. No samples were removed. Information on the donor was used as a covariate, using the design formula: “~patient + condition”.

#### Macrophages infected with Tuberculosis from (53)

We calculated differential expression between monocyte-derived macrophages from healthy donors infected with tuberculosis versus control samples. Datasets GSE133145 and GSE143731 were analyzed separately, but with a common design formula. Although there were also multiple treatments, the expression variance was almost exclusively driven by the difference in disease status. We therefore added the treatment as a covariate to the design formula (“~patient + treatment + condition”), but only investigated differential expression between infected macrophages and controls. One control and one infected sample from the GSE143731 series were removed from the analysis, as they were clear outliers in the PCA plot.

#### CD4+ follicular T-cells resting vs LPS-stimulated

We quantified differential expression between CD4-positive follicularT-cells in resting vs stimulated condition (47). The design formula used in DESeq is “~condition”.

#### AML subtypes

Differential expression was calculated using data from (90) and comparing samples with high leukaemia stem cell burden (GPR56-high) vs low leukaemia stem cell burden (GPR56-low samples) based on immunophenotyping as defined in (90). The design formula was: “~GPR56status”. We did not use shrunken log fold-changes as input for GRaNPA but we verified that results are qualitatively unchanged when doing so.

#### Tumour-associated and tissue resident macrophages from human breast tissue

Raw RNA-sequencing data was obtained from GSE117970 (55), and processed in the same way as described for expression data for TF K/Os. We obtained differentially expressed genes between tissue resident and tumour associated macrophages using the design formula “~condition”.

### GRaNIE: Construction of eGRNs

The following is needed as input for GRaNIE:

1. Raw or pre-normalized chromatin accessibility data (e.g., ATAC-seq, DNase-Seq or histone modification ChIP-seq data such as H3K27ac)
2. Raw or pre-normalized RNA-seq counts
3. Pre-compiled lists of TFBS predictions per TF (we provide predictions for human and mouse TFBS that were derived as described in (27))
4. TAD domains (optional)

For all datasets in this study, we used the same default parameters when constructing the eGRNs as described below.

GRaNIE is conceptually based on the procedure described in (7) and has the following main steps:

1. *Process chromatin accessibility and RNA-seq data* Both ATAC-seq and RNA-seq may be raw counts or pre-normalized counts. If raw counts are provided for RNA-seq, by default we quantile normalized the RNA-Seq count data in order to minimize the effects of outlier values that may otherwise have a large influence on the resulting correlations. For chromatin accessibility data, we employ a size factor normalization as implemented in DESeq2 (43). However, the user can define which type of normalization shall be used for either data. Additional filters for excluding particular chromosomes (e.g., sex chromosomes) or genes / peaks with low counts can optionally be used. The latter is implemented by removing genes / peaks if the average counts across all samples are below a specified threshold (5 by default).
2. *Overlap TF binding sites with ATAC-Seq peaks* Based on the provided list of putative TFBS per TF (see (27) for details), we overlap all TFBS from all TF with the open chromatin peaks and record for each peak and TF whether at least one putative TFBS is located within the peak. This binary TF-peak binding matrix is used in subsequent steps.
3. *Identify statistically significant TF - peak connections* To identify statistically significant TF-peak connections we implement a cell-type specific data-driven approach. In brief, we first calculate the Pearson correlation coefficients between the expression level of each TF and the open chromatin signal of each peak across all samples. We then use an empirical FDR procedure to identify statistically significant TF-peak connections. For this, for each TF, we split the peaks into two sets: A foreground set containing the peaks with a putative TFBS and a corresponding background set consisting of peaks without putative TFBS based on the TF-peak binding matrix calculated above. We then discretize the TF-peak correlation *r* into 40 bins in steps of 0.05 ranging from −1, −0.95, …, 0, …, 1 and calculate a bin-specific FDR value using two different directions (*positive*: left to right from −1 to 1, *negative*: right to left from 1 to −1). For each bin (correlation threshold) *k*, we calculate the empirical FDR according to the formula 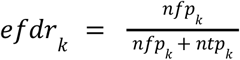, with *nfp_k_* and *ntp_k_* denoting the total number of TF-peaks in the background and foreground, respectively, for which *r* ≥ *k* (direction positive) and *r* < *k* (direction negative). To make the numbers from foreground and background compatible, we normalize *nfp_k_* beforehand by their ratio (i.e., 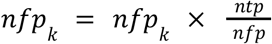, with *ntp* and *nfp* denoting the total number of TF-peaks in the foreground and background, respectively.
4. *Activator-repressor TF classification (optional)* Optionally, the TF classification as described in (27) can be run and is fully integrated in GRaNIE. It produces a classification of TFs into putative activators, repressors, or undetermined. Briefly, it compares the distribution of correlations for peaks with putative binding sites (foreground) against all other peaks (i.e., background) and classifies TFs depending on whether the correlations of putative targets are significantly more positive than (activator), more negative than (repressor), or indistinguishable from (undetermined) the background.
5. *Identify statistically significant peak-gene connections* Next, we add peak-gene connections to our network. We identify highly correlated peak-gene pairs based on their Pearson correlation and the associated p-value (using contest in R) between the normalized RNA-seq for the expression of a gene and the corresponding open chromatin peak. GRaNIE offers two options to decide which peak-gene pairs to test for correlation: in absence of additional topologically associating domain (TADs) data from Hi-C or similar approaches it uses a local neighborhood-based approach with a custom neighborhood size (default: 250kb up- and downstream of the peak) to select peak-gene pairs to test. In the presence of TAD data, only peak-gene pairs within a TAD are tested. The user has furthermore the choice to specify whether overlapping TADs should be merged or not. We offer two options of where in the gene the overlap with the extended peak may occur: at the 5’ end of the gene (the default) or anywhere in the gene. GRaNIE also records additional properties for each peak-gene pair such as their distance as well as gene type & status as annotated by Gencode. By default, only protein-coding and lincRNA genes are kept in the eGRN, but this can be customized to include other gene types.

### Quality controls

We implemented a range of quality controls for the different steps to assess signal vs noise in our eGRN. The package optionally offers PCA plots for both the RNA-seq as well as open chromatin data, and upon availability of additional metadata that can be provided, the PCA data can also be colored accordingly. This facilitates the detection of batch effects and outlier samples that may introduce unwanted variation.

For assessing TF-enhancer links, as described in the main text, we employ a permutation-based approach in addition to the real data for comparison by fully randomizing the TF-peak matrix.

For assessing enhancer-gene links we base our quality controls on the assumption that peak accessibility and gene expression is positively correlated. Thus assessing the amount of signal (i.e. small p-values) of positive correlations vs negative correlations serves as a proxy for signal to noise. We then assess the signal to noise in our real network vs a set of random peak-gene links. Specifically, we expect that positive correlations outnumber negative in the real network, and not in the randomized network. We also expect that with increasing distance the fraction of pos vs neg correlations gets smaller.

Lastly, we also perform the same procedure for randomized peak-gene combinations to empirically model the background distribution. For this, after selecting all peak-gene pairs, the sample IDs from the RNA-Seq data and the peak IDs from ATAC-seq are shuffled and the same procedure as described above is run. We include a number of QC plots as part of the output.

(6) Filter GRN connections and calculate peak-gene FDR

Lastly, we offer a variety of options to combine and filter TF-peaks and peak-genes to derive the full GRN for subsequent analyses. For example, both types of connections can be filtered by their FDRs or by their correlation, peak-gene links can further be filtered by their distance, gene type, and other criteria. By default, we retain only peak-gene pairs that are positively correlated. After applying all filters for the peak-gene links, multiple testing adjustment is performed using Benjamini-Hochberg. The default thresholds for TF-peak and peak-gene links are FDR < 0.2 and FDR <0.1, respectively.

Lastly, we provide heatmaps and boxplots that compare the connectivity for the real and random eGRNs.

### Molecular evaluation of GRaNIE TF-peak links using ChIP-seq

We obtained macrophage-specific ChIP-seq data from ReMap 2022 (92) for all TFs that were present in any of the GRaNIE inferred eGRNs (CEBPA, CEBPB, FOS, GABPA, GFI1, IRF8, IRF9, LYL1, MYB, NR1H3, PPARG, RUNX1, STAT1, STAT2, VDR). For these TFs we determined the overlap of GRaNIE inferred TF-linked peaks (independent of whether they also are linked to a gene) with the respective ChIP-seq peaks (within 10kb) and calculated the enrichment over the background set of ATAC-seq peaks that just contained the TF motif using Fisher’s exact test. We excluded two TFs (SPI1 and CTCF) that had more than 10k connections in GRaNIE at 0.4 or 0.5 FDR thresholds but no connections at lower FDR thresholds.

### Molecular evaluation of GRaNIE peak-gene links using eQTL data

We downloaded c/s-eQTL data from the eQTL catalogue (https://www.ebi.ac.uk/eqtl/, accessed on May 5th 2022), selecting all six datasets with eQTLs in monocytes or macrophages. We combined the eQTLs from all datasets and filtered the associations to have a permutation-based FDR < 0.3. Only GRN peaks that harbour at least one eQTL SNP in these datasets can be evaluated. For every peak, we overlapped the eQTL SNPs, and counted a peak-gene link as validated if any eQTL SNP affected the same gene as present in the GRN. To test whether the overlap between peak-gene links and eQTL is enriched as compared to a random background, we also validated links between the GRN peaks and randomly sampled distance-matched genes based on 50kb bins. We repeated the random background sampling 20 times and calculated the odds ratio between validated GRN links and validated background links. We calculated the enrichment of validated links by eQTL overlap for a range of peak-gene FDR thresholds in each of the four macrophage GRNs.

### Molecular evaluation of GRaNIE TF-peak-gene links

We obtained the tracks of human enhancers identified using CAGE data from FANTOM5 that were differentially expressed in T-cells and macrophages (30). Track coordinates were translated to GRCh38 using UCSC *liftOver* tool (93). We quantified the overlap of GRaNIE eGRN enhancers (peaks linked to a TF and a gene) and the respective cell-type specific (macrophage in **Fig. 1F,2H** and T-cell in **Fig. 2H)** CAGE peaks and tested for the association of eGRN peaks (vs all other peaks within 250kb of a TSS) and cell-type specific CAGE enhancers using Fisher’s exact test.

### Utility evaluation of GRaNIE eGRN regulons using GO-term specificity

To assess the specificity of the top GO terms enriched for the predictive TFs **[Fig. S9A],** we randomised the TF-gene connections and recalculated the enrichment. Manual inspection suggested that the returned terms for the random network are more general and often unrelated to macrophage biology. To quantify the specificity, we plotted the distribution of the number of genes annotated to each returned GO term for the real network, and for five permuted networks **[Fig. S9B],** following the rationale that more general terms would have more genes associated with them in the database.

### Downstream analyses implemented in GRaNIE

The following functionalities are available within the GRaNIE package:

- Descriptive statistics pertaining to the structure of the graph, such as the number of nodes and edges and their types, the distribution of node degrees, and the top nodes with regard to degree centrality and eigenvector centrality.
- Community identification, for which the package supports multiple algorithms (louvain, walktrap, leading eigenvector, fast greedy, and optimal). The communities can be supplemented with descriptive statistics similar to those previously described, but specific to each individual community.
- Enrichment analyses in three different flavours; a general enrichment analysis for the whole network, a community-based enrichment analysis, or a TF-based enrichment analysis.

### GRaNPA: Prediction model

For the prediction model, first we need to compute the differential expression resulting from a perturbation between two different conditions, for which we used DESeq2. We define DE(j) as the differential expression value for the j^th^ Gene.

Second, we need to define the GRN as a mathematical function:

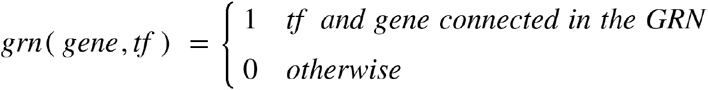

Based on this, we construct matrix *X*, a relation matrix between genes and TFs where each row is a gene and each column is a TFs. There is a 1 in the cell for a gene and TF if and only if they are connected through the GRN.

Finally, we use Random Forest (RF) Regression based on the following formula to predict a differential expression value for each gene *i* based on its connected TFs *X_i,tf_*:

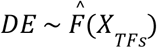

RF has been implemented using the “ranger” package in R. To avoid overfitting we use 10-fold cross validation; no hyper-parameter tuning on the random forest was performed. We measure performance by assessing the cross-validation R^2^.

### GRaNPA: Construction of the randomized network to assess edge-specificity

To assess the edge-specificity of our network during the random forest regression we constructed a randomized control network based on the structure of the actual GRN. It has the same number of edges with the same degree distribution for TFs with only the gene labels permutated so the connection between TFs and genes are randomized. The differential expression values are unchanged. In case we use any weighing method for the edges, the same method will be applied for each edge of the randomized version to recalculate weight if needed.

### GRaNPA: Random signal generation to assess overfitting

To control for over-fitting in the random forest regression, we used the real network structure and assigned randomly generated values to genes instead of their differential expression value. The random values are chosen from a uniform distribution with minimum and maximum of −5 and 5 respectively. Any prediction value for this network is a consequence of overfitting. The reason we used a uniform distribution is that we want to check overfitting and we don’t want the distribution to be like any differential expression distribution.

### Benchmarking against other networks / tools

#### Dorothea

Dorothea [11] is a resource containing TF-target interactions. The connections are tagged by a confidence level based on the number of supporting evidence. The confidence levels are from A considered to be highly reliable to B-D reserved for curated and/or ChIP-seq interactions and the lowest confidence level E, for connection with only computational support.

#### TRRUST

TRRUST (16) is a database of transcriptional regulatory networks for humans and mice. It has been constructed using text-mining followed by manual curation. TRRUST v2 regulatory network for humans contains 795 TFs, 2067 genes and 8427 regulatory links. We did not use any weighing for the edges as it was not provided by TRRUST v2.

#### ChEA3

ChEA3 (17) are TF-target gene libraries containing targets determined by ChIP-seq experiments from ENCODE, ReMap and publications. It also contains co-expression connections based on RNA-seq data from resources like GREx and ARCHs4.

#### ANANSE

ANANSE (5) is enhancer-based cell type specific network that can predict key transcription factors in cell fate determination. We used ANANSE macrophage specific network filtered by 0.8 probability for its links.

### Assessing the cell-type specificity of network models

In order to assess cell-type specificity of network models, we check their performance on different DE data from other cell-types. In this analysis, we used naive macrophages, naive T-cell and AML and evaluated their performance for DE data from GPR56-positive vs GPR56-negative AML, resting vs stimulated CD4 positive follicular T-cell (in 10 different subcell-types) and naive-macrophages vs 5-h infected with Salmonella. We filtered genes using 0.1 adjusted p-value and 1 absolute log fold change thresholds.

### Enrichment of regulons among TF K/O data

We obtained cell-type specific knock-out (K/O) data for THP1 derived macrophages (NFKB1) (49), the human AML cell line MV4-11 (IRF8) (50), and primary human CD4+ T cells (IRF1 and IRF2) (44). Differential expression analysis was performed using DESeq2 (43), comparing all K/O conditions to their non-targeting controls. All genes significantly downregulated upon TF K/O (adjusted p-value < 0.05 and fold-change < 0), were considered as TF targets. Enrichment analysis of TF targets in the cell type specific regulons from the macrophage, AML and T-cell eGRNs was performed using Fisher’s exact test as implemented in the *GeneOverlap* package in R. Enrichments with at least five overlapping genes and Fisher’s exact p-value <0.05 were considered significant.

### Calculating variable importance measures

We used the permutation approach to measure variable importance using the “ranger” package in R. This accuracy-based approach uses the out-of-bag sample to calculate the importance of a specific variable. The importance is based on the difference in the prediction accuracy of out-of-bag sample and the prediction accuracy of out-of-bag sample while its variables have been randomly shuffled while all other variables kept the same.

### Visualization (Shiny App)

We provide a web application based on a Shiny App for interactive visualization of the eGRNs for different cell types (https://apps.embl.de/grn/).

### Gene set enrichment analysis

Pre-ranked gene set enrichment analysis (GSEA, ranking based on log2 fold-change) was performed using the Bioconductor/R package *fgsea* (94). The M1 macrophage signature gene set was obtained from (95).

### GWAS enrichment

We tested if the macrophage GRNs were enriched for genetic heritability of GWAS traits using stratified linkage disequilibrium score regression (S-LDSC) (67). 806 GWAS summary statistics that included participants of European descent were downloaded from the GWAS *Catalog* (96), harmonized and converted to the LDSC format as described on the LDSC Github repository (https://github.com/bulik/ldsc). We removed summary statistics with fewer than 10,000 individuals and fewer than 100,000 SNPs, because those studies were likely underpowered, leaving 442 traits. We created peak sets for each of the three macrophage GRNs by extracting the enhancer regions that were present in the GRNs after filtering for peak-gene distance, TF-peak FDR and gene differential expression effect size. We used 54 sets of general genomic features (downloaded from https://alkesgroup.broadinstitute.org/LDSCORE/, following (97)) and a peak set based on all macrophage enhancers within 250kb of genes as background regions. Adding these regions as a background ensures that the identified enrichments are not due to the general enrichment of heritability in (macrophage) enhancers near genes, but specifically in the enhancers that are part of the GRNs. We calculated the heritability enrichment p-values and corrected them for multiple testing within each trait.

### GO enrichment of GRNs

The general enrichment analysis was run using *topGO* (v2.42.0), with the foreground being the genes in the filtered GRN, and the background being the genes within a predefined 250kb neighborhood of the peaks in the GRN. In more specific enrichment analyses such as those for the top transcription factors, which are ranked by their predictive capacity, the foreground is selected based on the genes a given TF is connected to within the network. Similarly, in community-based enrichment analyses, the foreground is simply the genes that are classified to a given community. To calculate the enrichment, a Fisher test is used alongside the weightOI algorithm, which is a mixture of the *“elim”* and the *“weight”* algorithms introduced by (98), to account for the GO hierarchy. Additionally, terms with less than 4 significantly annotated genes were omitted from the results in the figures. For the sake of better visualizing the enriched terms, the figures are limited to the top ten enriched terms per category. The full list of enriched terms can be found in **Tables S7 and S8.**

### Identifying targets for fine-mapped GWAS-SNPs using eGRNs

Fine-mapped GWAS variants for autoimmune diseases were generated using probabilistic identification of causal SNPs (PICS) algorithm. These variants were downloaded from i) previously published list and lifted to the *hg38* build (99) and ii) data portal under the filename “PICS2-GWAScat-2020-05-22.txt.gz” from https://pics2.ucsf.edu for the hg38 build. We kept all variants with a PICS probability of greater than 50%. We identified their target genes by overlapping these variants with the peaks from different macrophage eGRNs and then using the peak-gene links from the respective eGRNs to assign the target genes **(Fig. 5C).** The full list can be found in **Table 3.** 24. Stone M, Li J, Grace McCalla S, Fotuhi Siahpirani A, Periyasamy V, Shin J, Roy S. Identifying strengths and weaknesses of methods for computational network inference from single cell RNA-seq data. bioRxiv 2021.06.01.446671;

