## Supplementary figures for "GRaNIE and GRaNPA: Inference and evaluation of enhancer-mediated gene regulatory networks applied to study macrophages"

### Supplementary Material

#### Supplementary Tables S1-10

**Supplementary Table S1-S6**: eGRNs for macrophages (S1-S4, naive - S1, primed - S2, infected- S3, union of naive and infected - S4) as well as for AML (S5) and CD4+ T-cell (S6) using the default parameters as described in the main text.

*Description of columns:* (1) *TF.name* - Name of TF, (2) *TF_peak.r* - Pearson correlation of TF and peak, (3) *TF_peak.fdr* - TF-peak FDR, (4) *peak.ID* - peak ID (coordinates), (5) *peak_gene.distance* - Genomic distance between peak and gene (in bp), (6) *peak_gene.r* - peak-gene Pearson correlation, (7) *peak_gene.p_raw* - peak-gene raw p-value, (8) *peak_gene.p_adj* - peak-gene adjusted p-value (FDR), (9) *gene.ENSEMBL* - Ensembl ID of gene, (10) *gene.name* - Name of gene. For the AML network in S5 coordinates are *hg19*, for all others *hg38.*

**Supplementary Table S7**: GO enrichments for the communities of the naive, primed and infected macrophage eGRNs.

*Description of columns: eGRN* - The name of the eGRN network, *community* - the ID of the community, *GO.ID* - GO ID, *Term* - the GO term as text, *Annotated* - number of genes associated with this term in total, *Significant* - the number of genes found in the foreground associated with this term in total, *Expected* - expected number of terms associated with this term given the foreground size, *pval*: raw p-value, *GeneRatio* - the ratio of *Significant* divided by the foreground size

**Supplementary Table S8**: GO enrichments for the important TFs for the naive, primed and infected macrophage eGRNs. Description of column same as Table S7.

**Supplementary Table S9**: List of GWAS traits used for the LDSC analysis. The summary statistics were from the GWAS catalog as well as from the LDSC website with pre-processed GWAS statistics.

**Supplementary Table S10**: References for enriched traits related to macrophage biology

#### Supplementary Figures 1-20

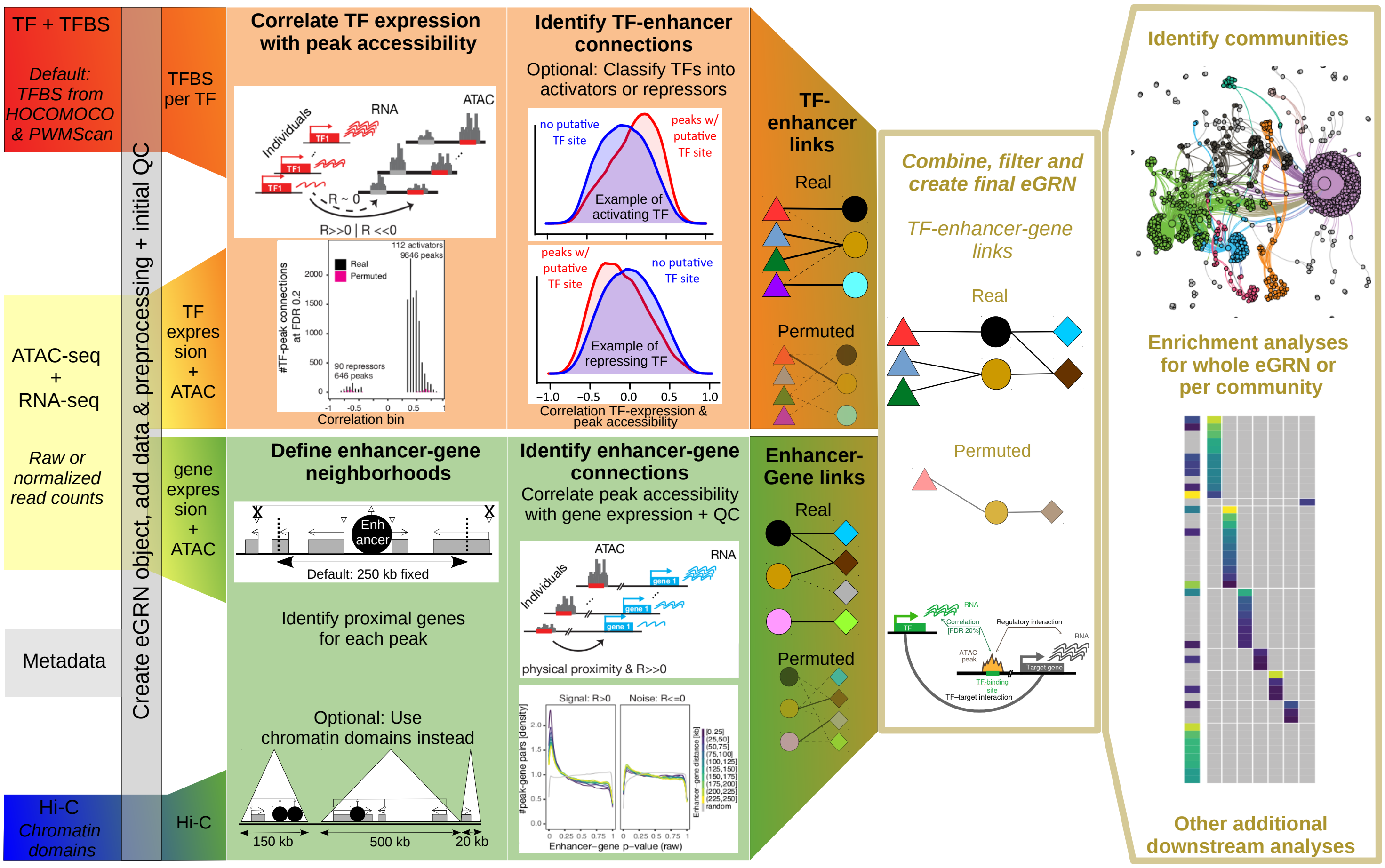


**Supplementary Figure S1**: Detailed graphical workflow of *GRaNIE*. Input for the package is highlighted at the left side, which includes mandatory and optional (Hi-C data) input. For methodological details, see the main text and Methods.

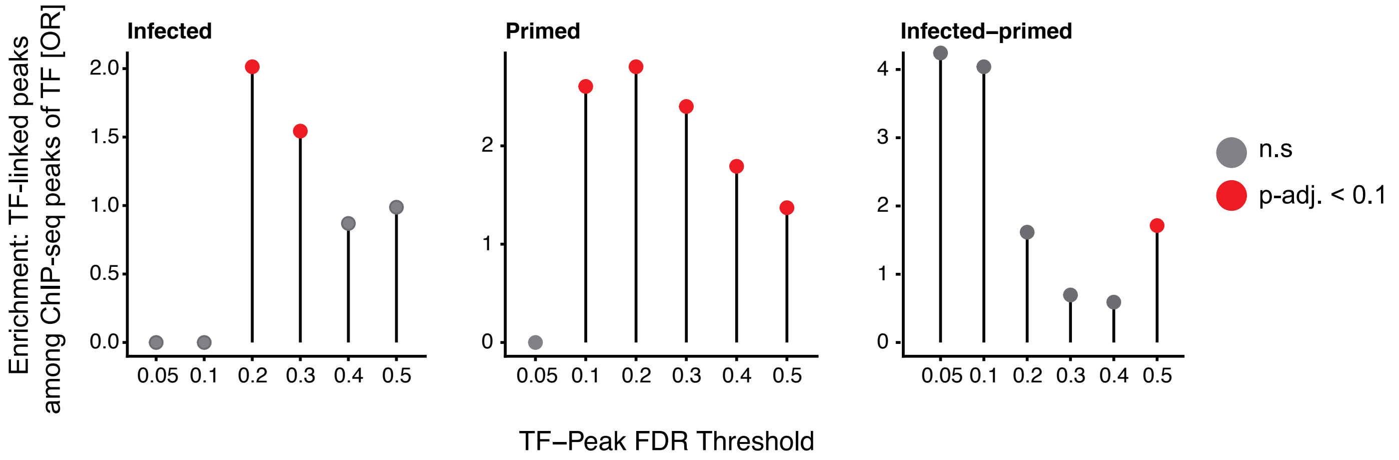


**<u>Supplementary Figure S2</u>**: Validation of the eGRN TF-peak links with ChIP-seq data. Enrichment of ChIP-seq peaks overlapping a GRaNIE-inferred TF-bound peak (same TF) are shown for different TF-peak FDRs in the infected (left), primed (middle) and primed-infected (right) macrophage eGRNs. Background: peaks that contain the motif for the respective TF but are not significantly linked.

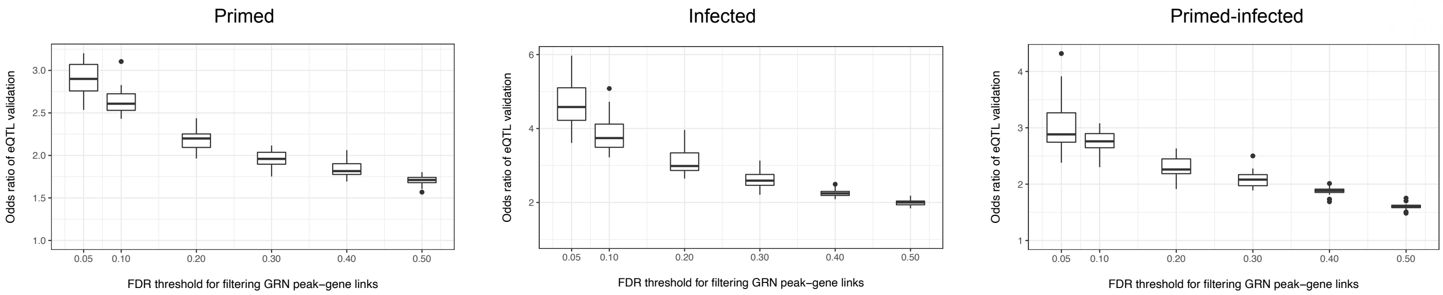


**<u>Supplementary Figure S3</u>**: Validation of the eGRN peak-gene links with macrophage eQTLs. Plots show the enrichment of eGRN links overlapping an eQTL over randomly sampled distance-matched peak-gene links overlapping an eQTL for different peak-gene FDRs in the primed (left), infected (middle) and primed-infected (right) macrophage eGRN.

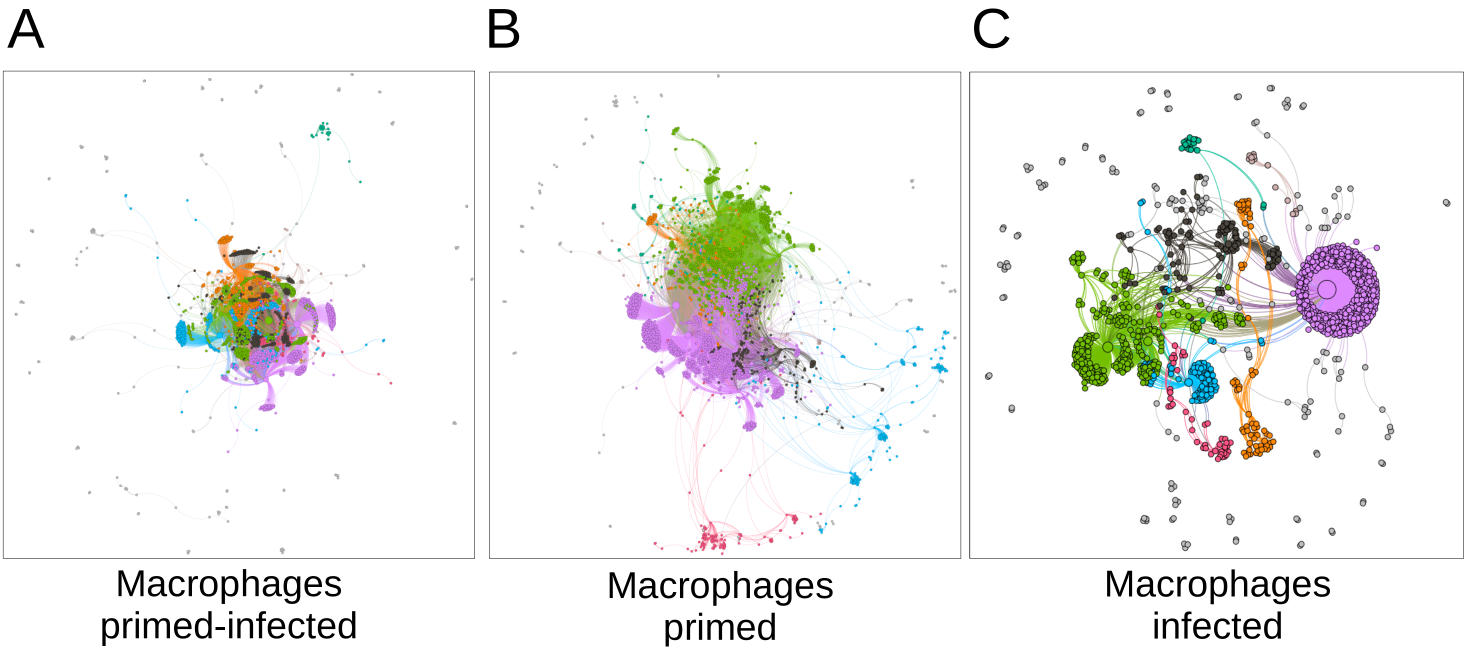


**Supplementary Figure S4**: Network visualizations and community identification of the other Macrophage eGRNs described in this manuscript in analogy to **Fig. 1E** using a forced-directed visualization. The colors correspond to the identified network communities.

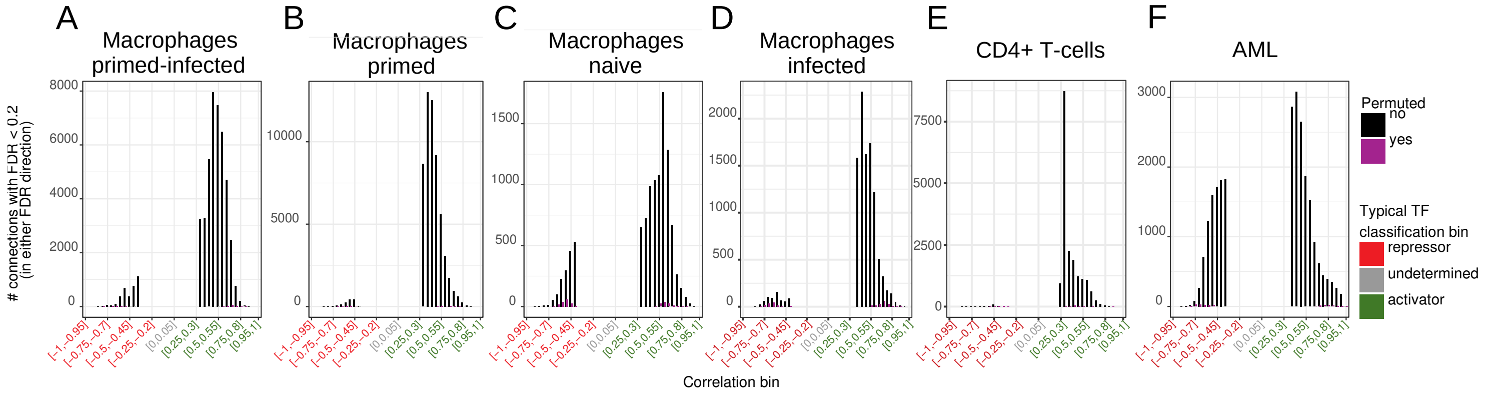


**Supplementary Figure S5**: QC and summary from the Macrophage eGRNs for the TF-peak connections. The histogram of the number of connections for which TF-peak FDR < 0.2 (y-axis), stratified by the TF-peak correlation bin (x-axis, in bins of 0.05) for real (black, labeled as ”no”) and permuted (violet, labeled as ”yes”) networks.

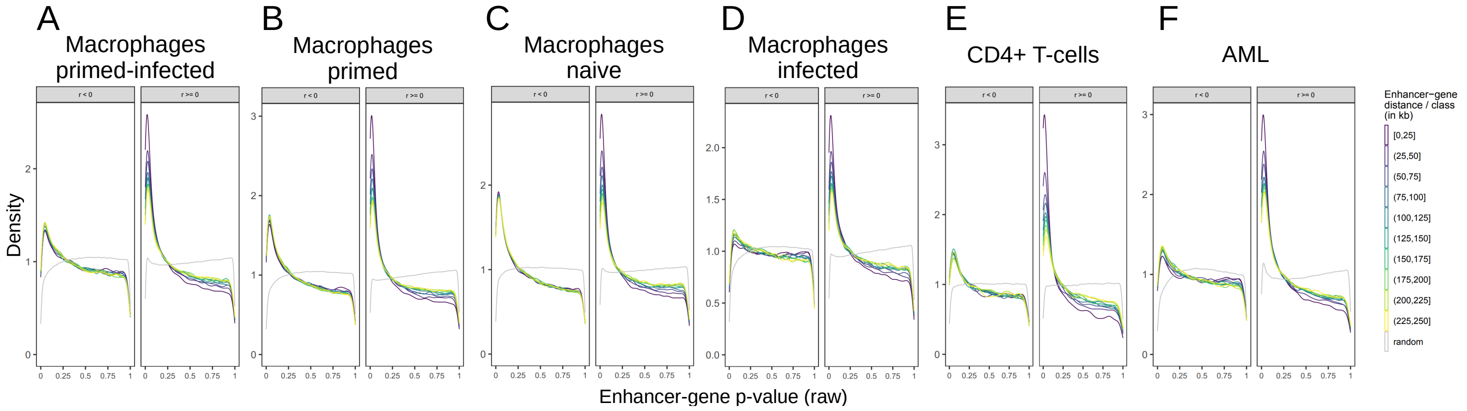


**Supplementary Figure S6:** QC and summary from the Macrophage eGRNs for the peak-gene connections. Distributions of p-values are shown for positive (right) and negative (left) peak-gene correlations. Peak-gene pairs are stratified by their distance (heat colors) and permuted gene-peak pairs (randomized network) are shown in gray.

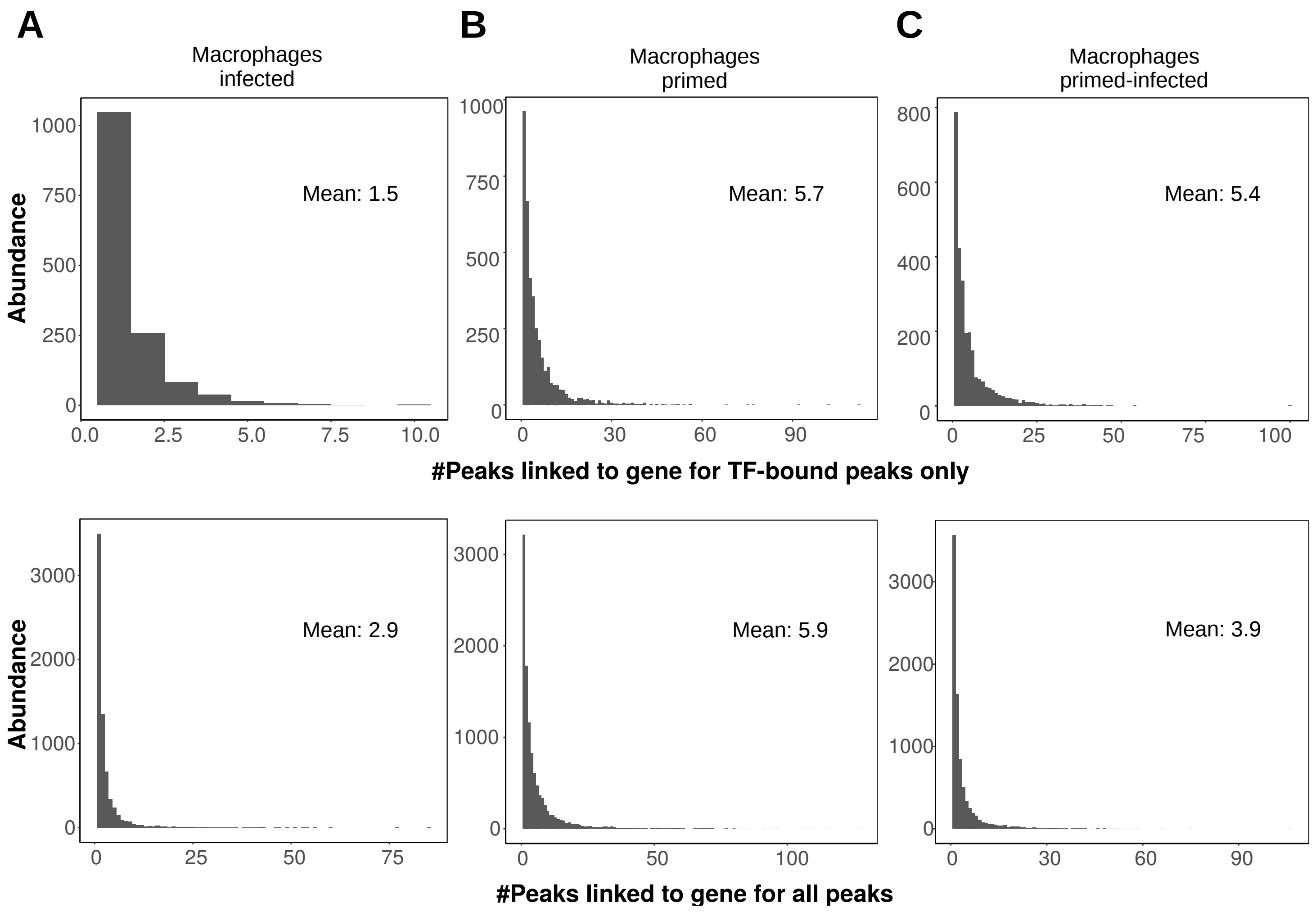


**Supplementary Figure S7:** Histograms of the number of peaks linked to a gene for the various macrophage eGRNs, along with their mean value. The upper row counts peaks only if they are TF-bound (i.e., as GRaNIE outputs them as proper TF-peak-gene connections), while the lower row includes all peaks (i.e., also those not connected to a TF, focusing therefore on all significant TF-gene connections and ignoring the TF-peak FDR).

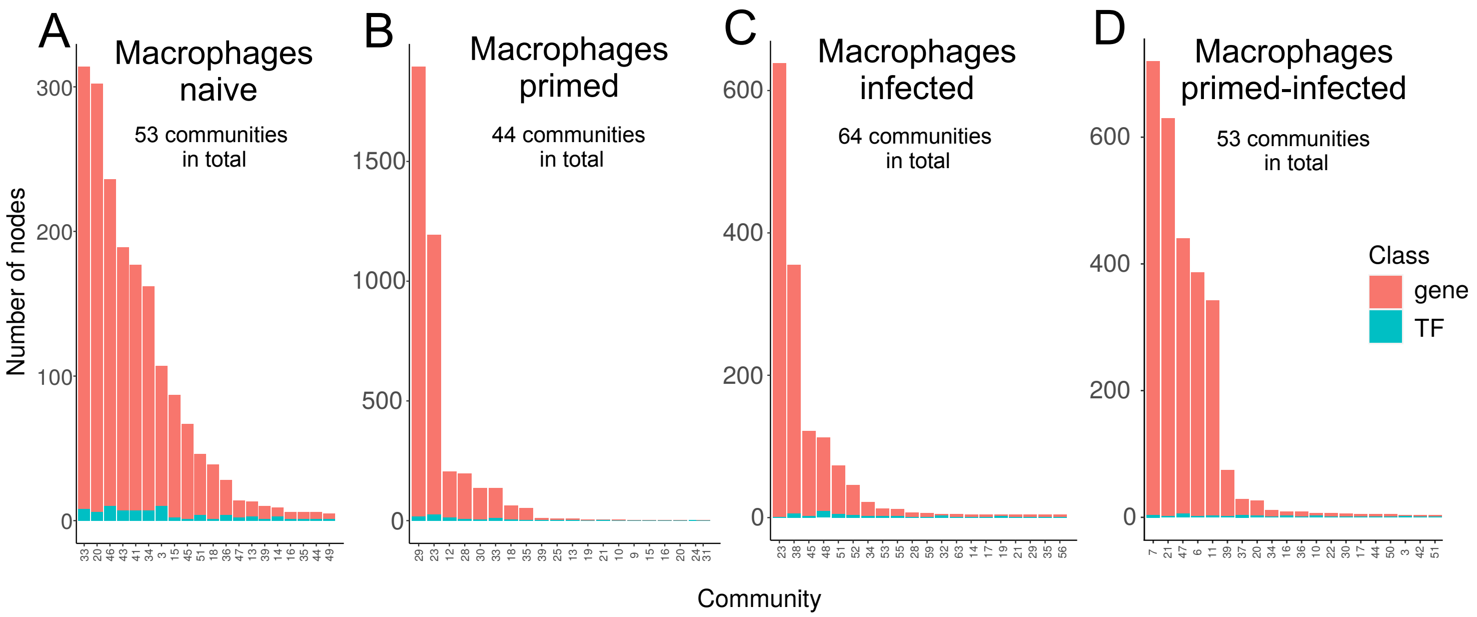


**Supplementary Figure S8.** Community sizes for the different macrophage eGRNs with respect to the number of genes and TFs per community.

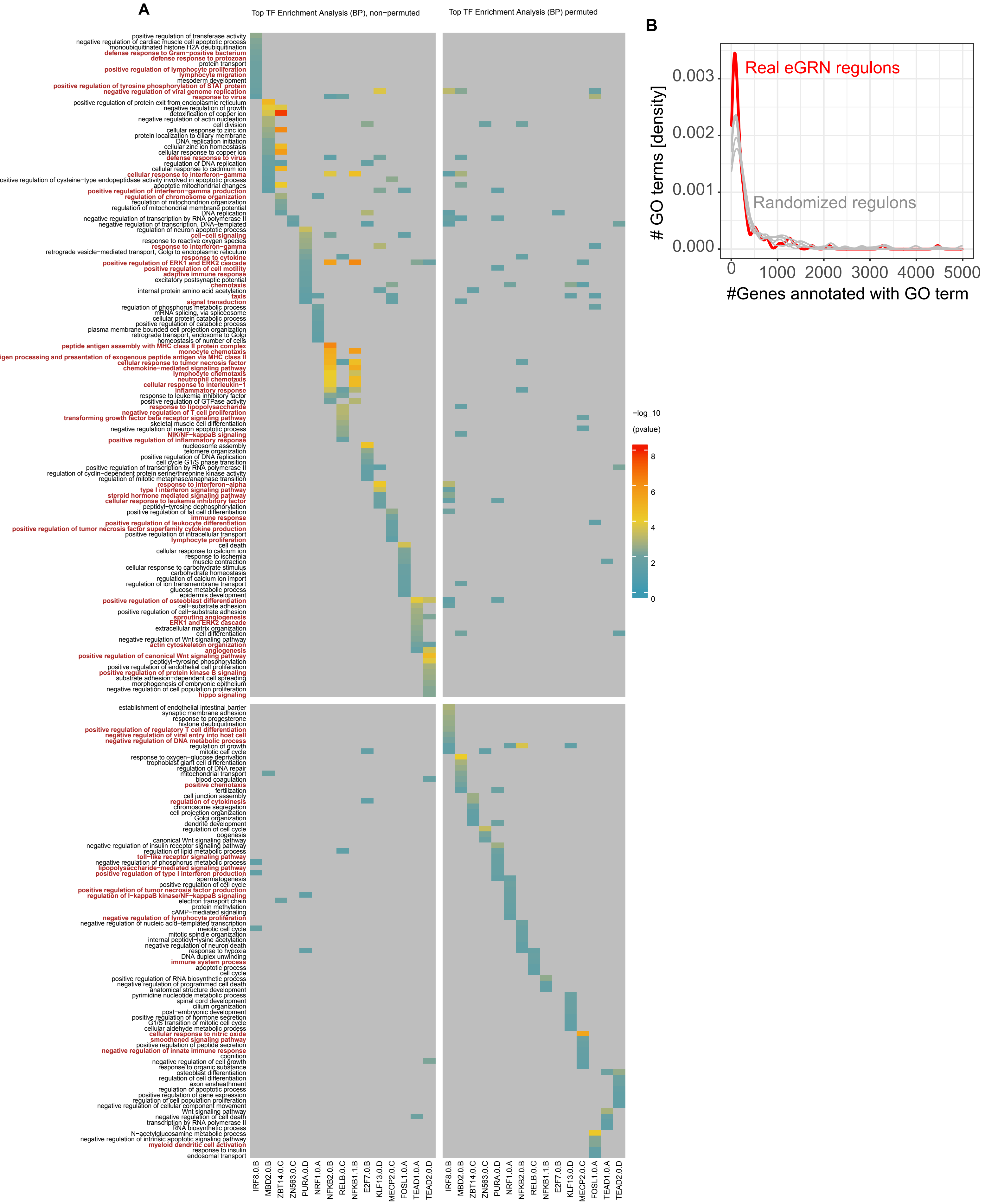


**<u>Supplementary Figure S9:</u> A)** Heatmap of GO enrichment for TF regulons of the real (union of the naive and infected macrophage eGRNs; left) and permuted TF-gene links (right). The real TF regulons have more macrophage related terms (manually labeld in red) than the permuted regulons. **B)** Distribution of the number of genes annotated to the top enriched GO terms in regulons of the real (red) and five permuted (grey) eGRNs. Less of annotated genes indicates higher specificity.

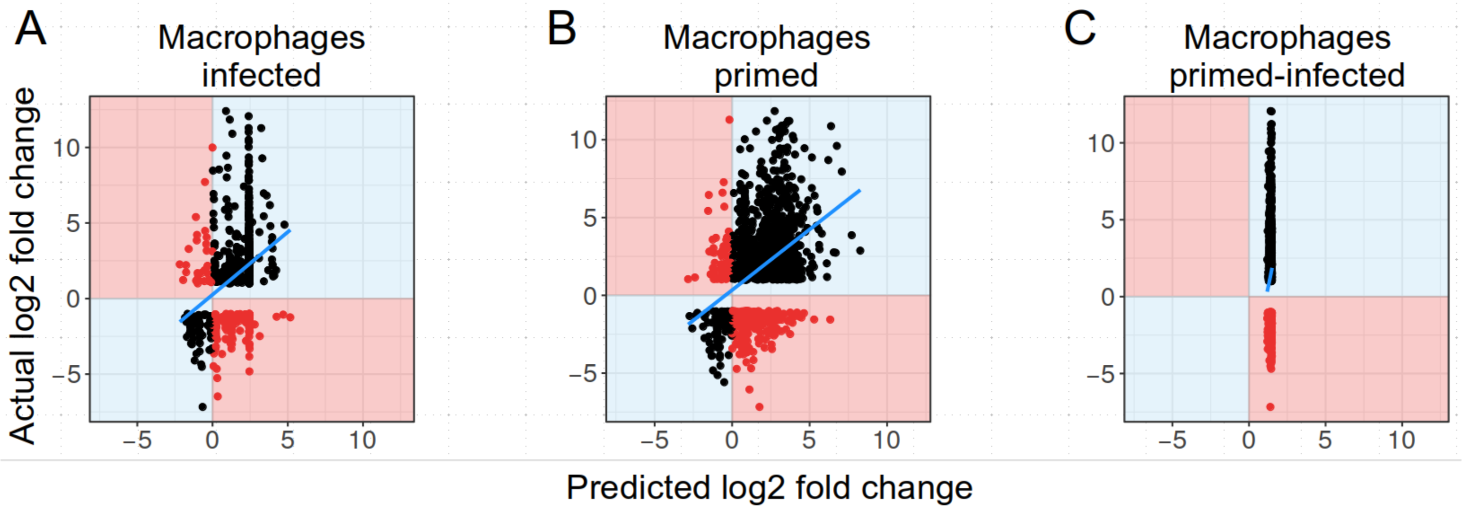


**Supplementary Figure S10.** True vs predicted (using GRaNPA) log2 fold changes for the macrophage expression response to salmonella infection are shown as scatter plot for three macrophage eGRNs, in analogy to **Fig. 2B**. For A and B, GRaNPA is able to predict the response, while this is not the case for C.

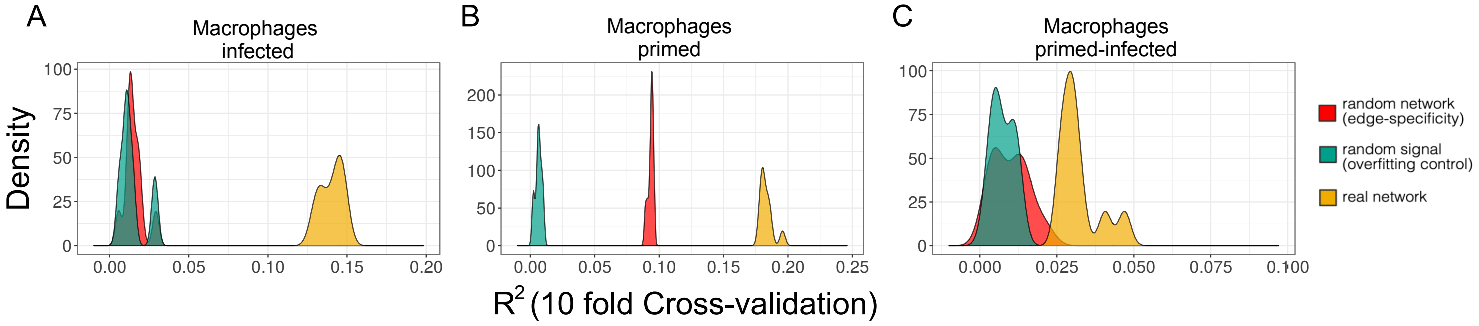


**Supplementary Figure S11:** Output of GRaNPA in analogy to **Fig. 2C** for infected (A), primed (B) and primed-infected (C) macrophage eGRNs. Distributions of $R^2$ for 10 random forest runs for predicting differential expression upon salmonella infection are shown as a density plot.

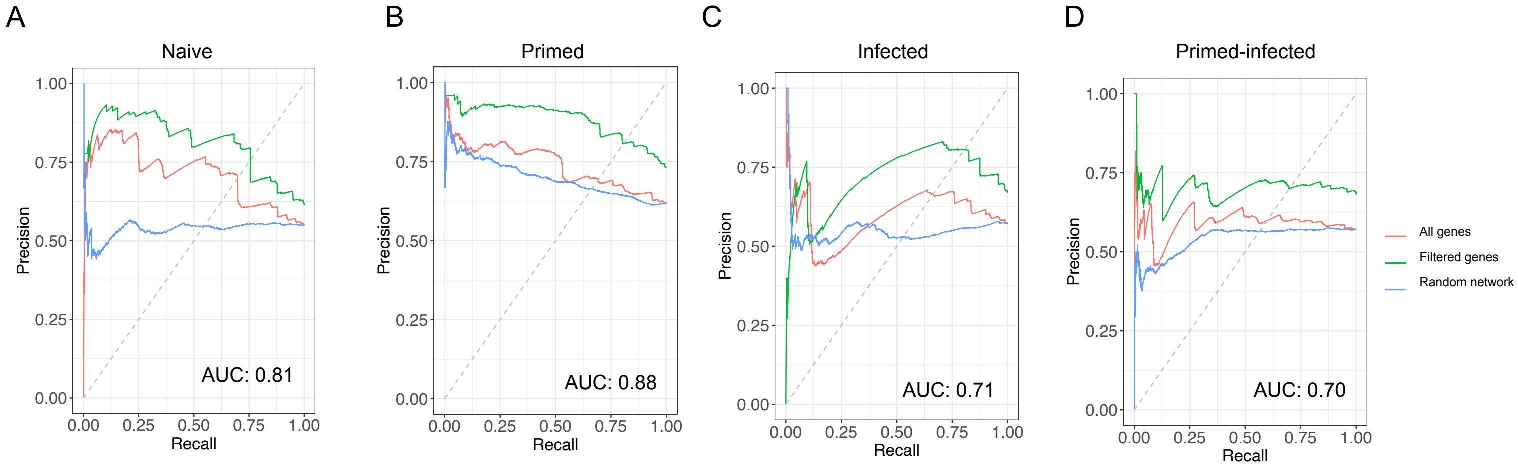


**Supplementary Figure S12:** Output of GRaNPA for classification of direction of differential expression data for different macrophage eGRNs (precision-recall curve). Random forest classification has been trained on the real network predicting all genes (red), the real network predicting genes with absolute log-fold change > 1 (green) and the random network predicting all genes (blue).

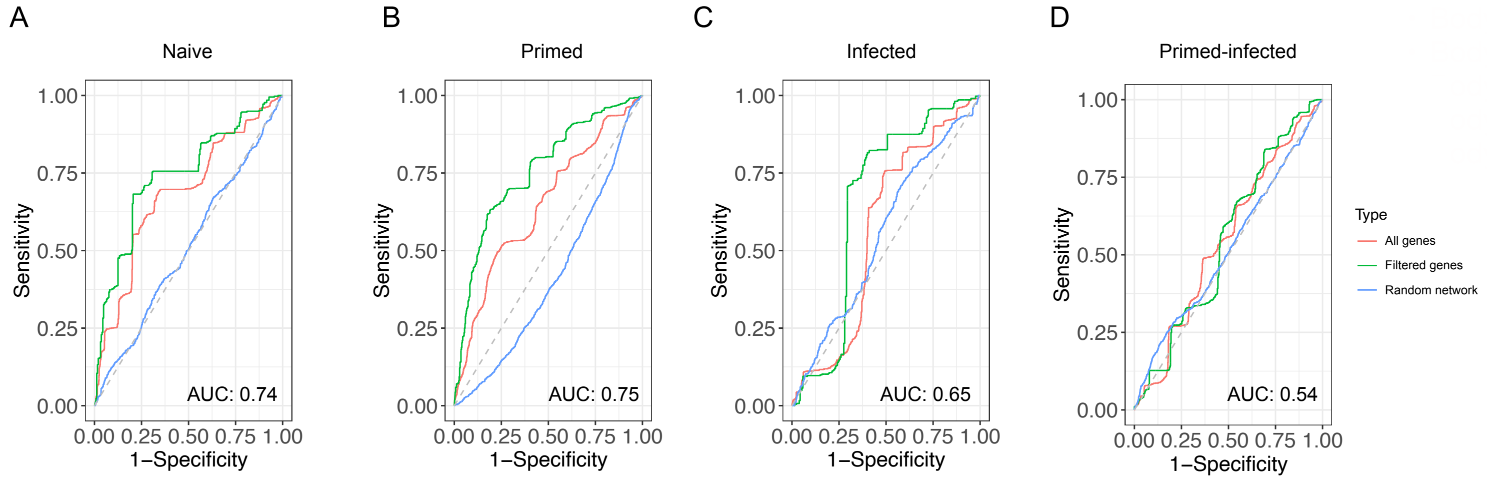


<u>**Supplementary Figure S13:**</u> Output of GRaNPA for classification of direction of differential expression data for different macrophage eGRNs (identical to **Fig. S12**, just showing the receiver operating characteristic - ROC - curve instead). For more details, see the caption for **Fig. S12**.

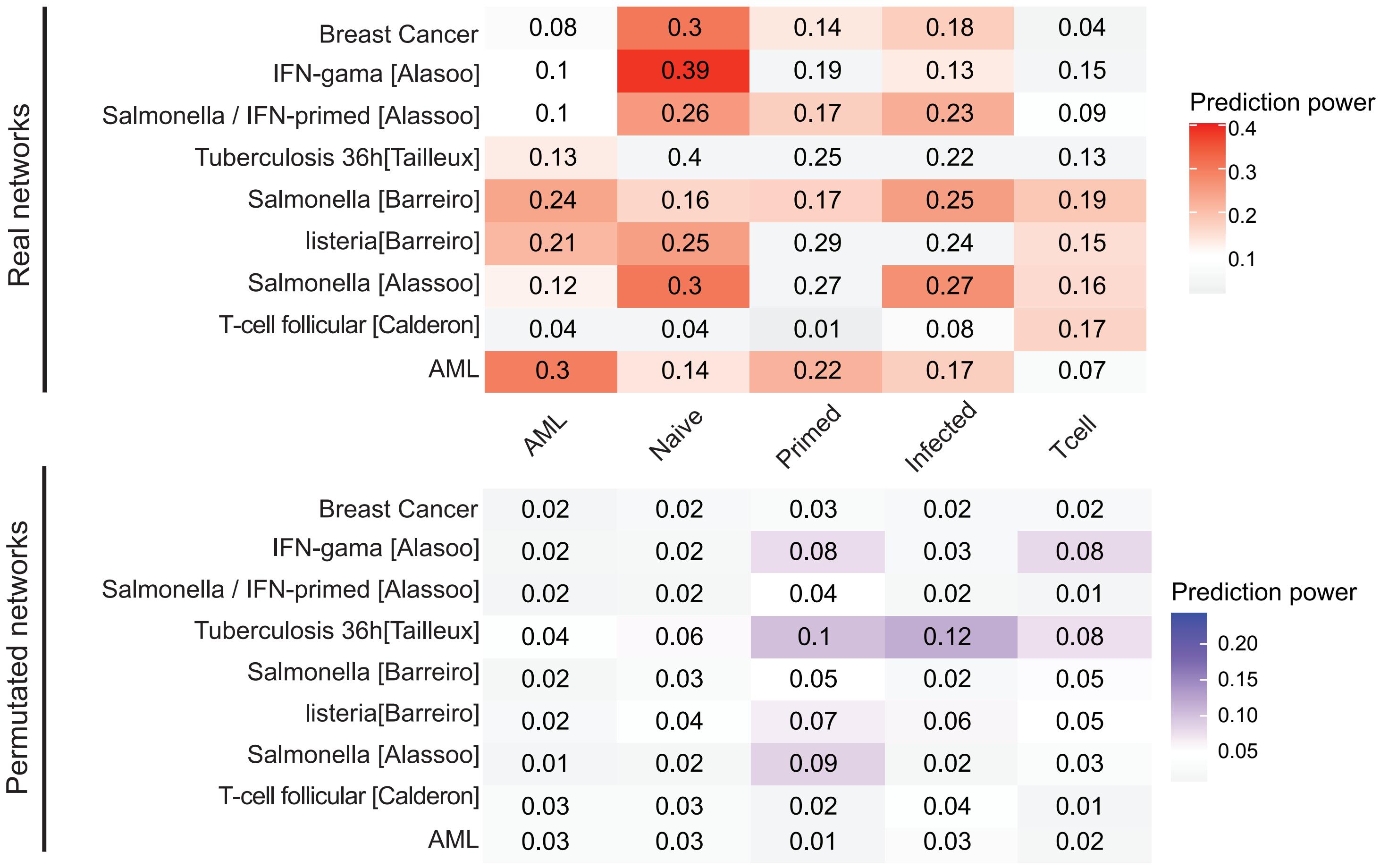


**<u>Supplementary Figure S14</u>:** Output of GRaNPA (mean R2 across 10 random forest runs) using a modified model where expression variation is included as gene-specific feature. Expression variation is calculated based on GTEX data. The R-squared values are shown for predicting the differential expression response in nine distinct perturbations (rows) with distinct GRaNIE-inferred eGRNs (columns; top) and with the respective random networks (columns, bottom). eGRNs that show a performance > 0.05 with the random eGRN (bottom) are colored in grey for the real network (top).

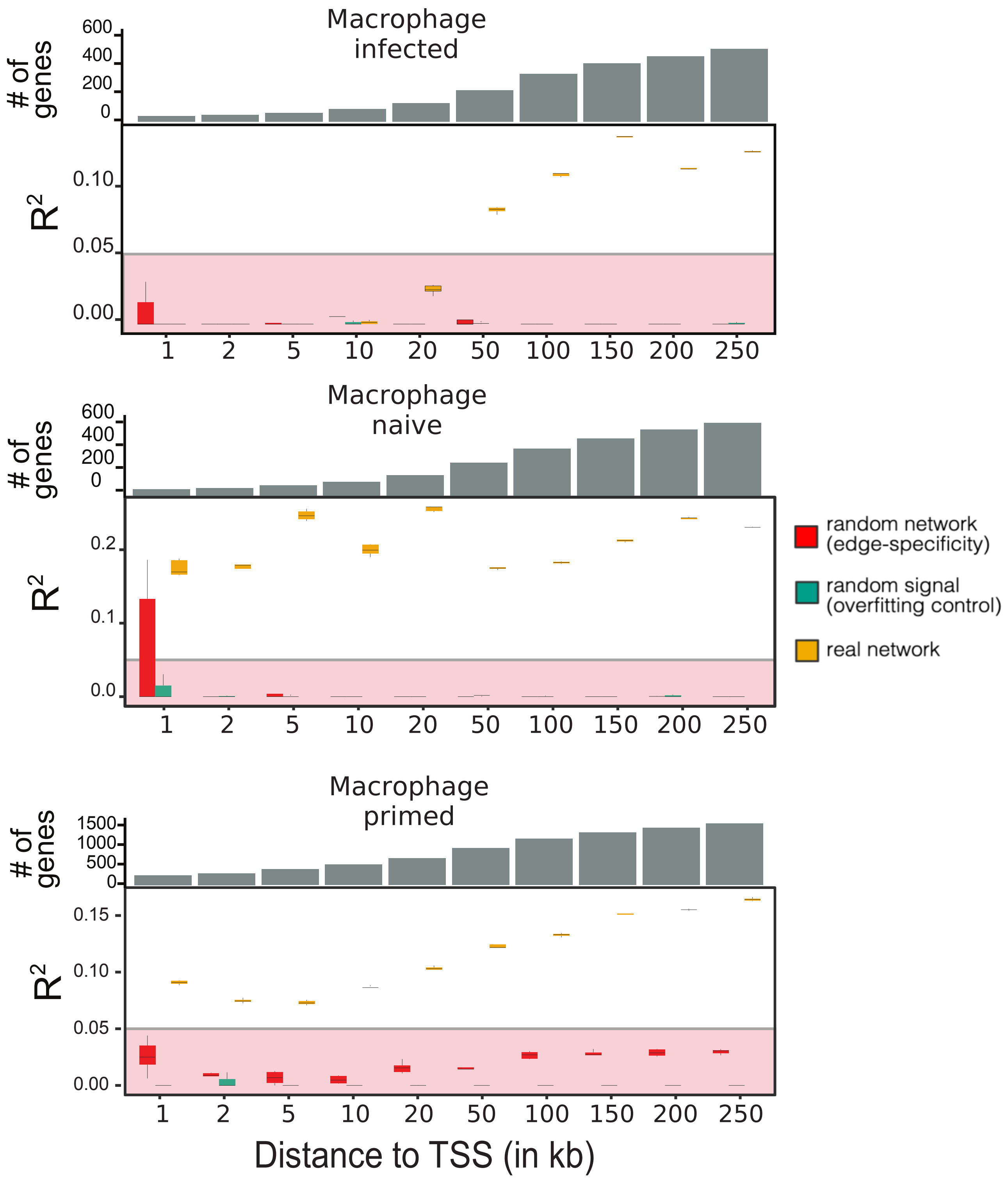


**Supplementary Figure S15 :** GRaNPA evaluation of subnetworks stratified by gene-enhancer distance for the eGRNs from infected, naive and primed macrophages (top to bottom) using differential expression between Salmonella infection after 5h and naive macrophages. For any particular distance threshold *k*, the subnetwork consists of all connections with a gene-enhancer distance of 0 to *k*.

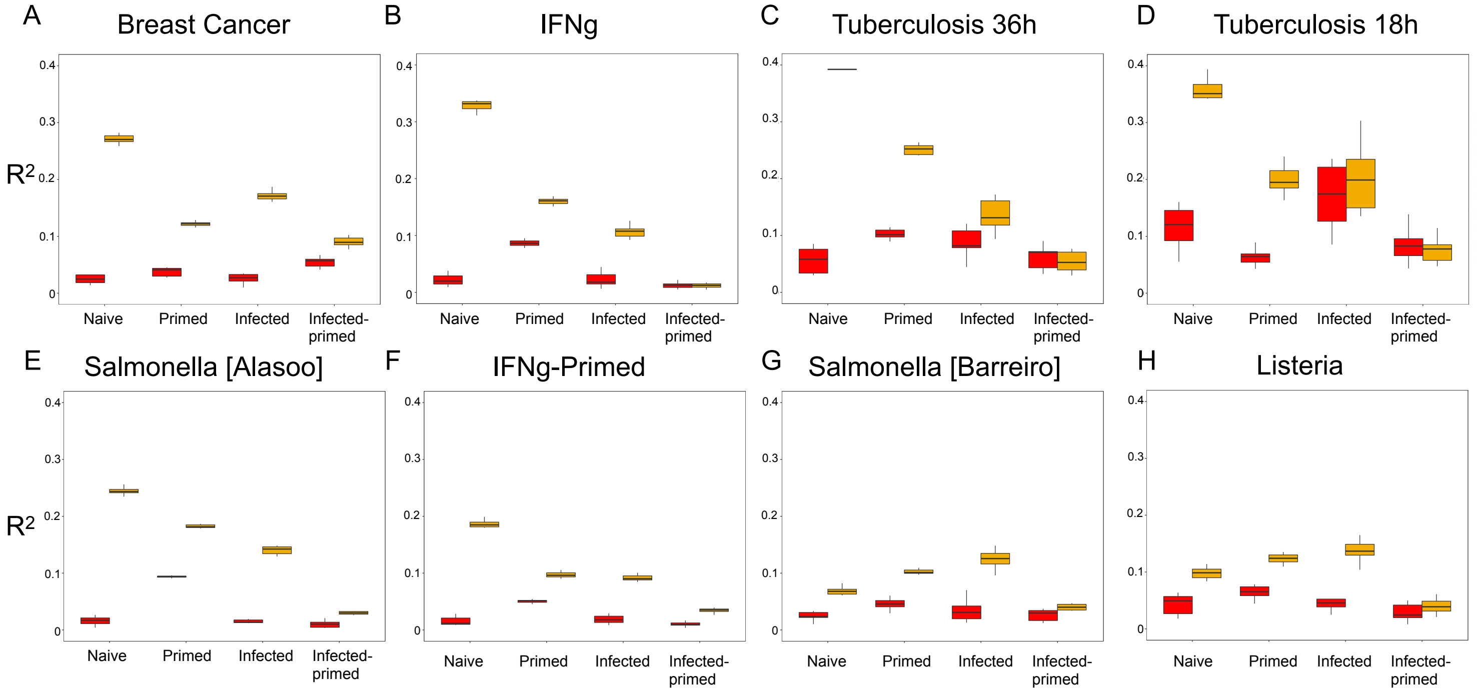


**Supplementary Figure S16:** GRaNPA evaluation of naive (left), primed (middle-left), infected (middle-right) and primed-infected (left) eGRNs across seven conditions of macrophage differential expression response. Overall, the naive and the naive-infected eGRNs were able to predict the differential expression response to most conditions ($R^2$>0.1 and $R^2$<0.05 for the respective random networks), while the primed network failed the specificity control for some. Since naive and infected eGRNs both showed high predictive power in these settings, we assessed the pathogen response using the union of the naive and infected macrophage eGRNs (**Fig. 4A**). GRaNPA evaluation of the primed-infected eGRN failed for most of the conditions, in line with its poor evaluation with respect to ChIP-seq data (**Fig 1C**).

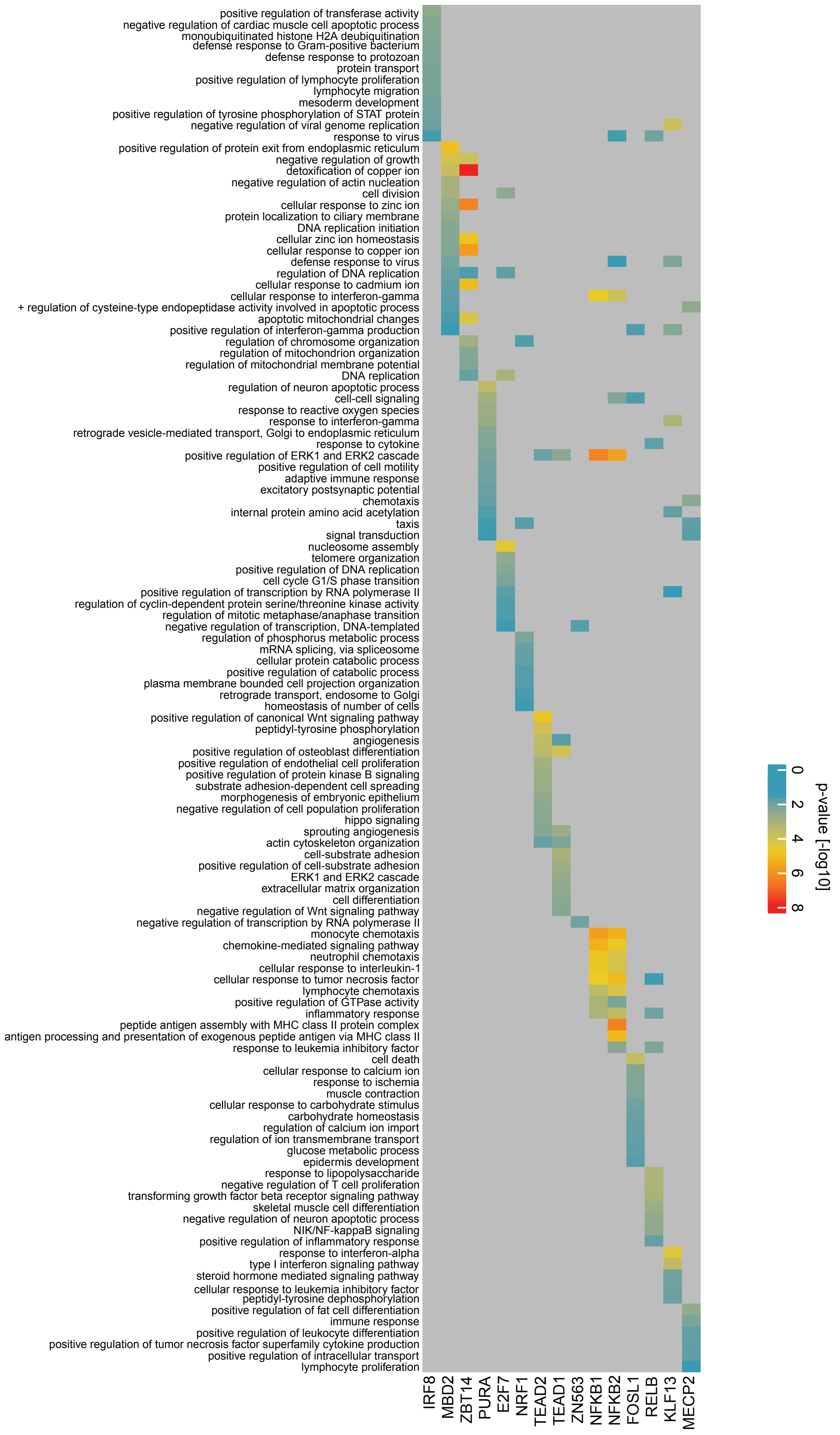


**Supplementary Figure S17:** GO enrichment analysis of the regulons of the 15 most important TFs across all conditions (as implemented in GRaNIE) in the union (naive+infected) macrophage eGRN revealed that each TF is associated with very specific terms, likely reflecting different defence mechanisms triggered by the pathogens.

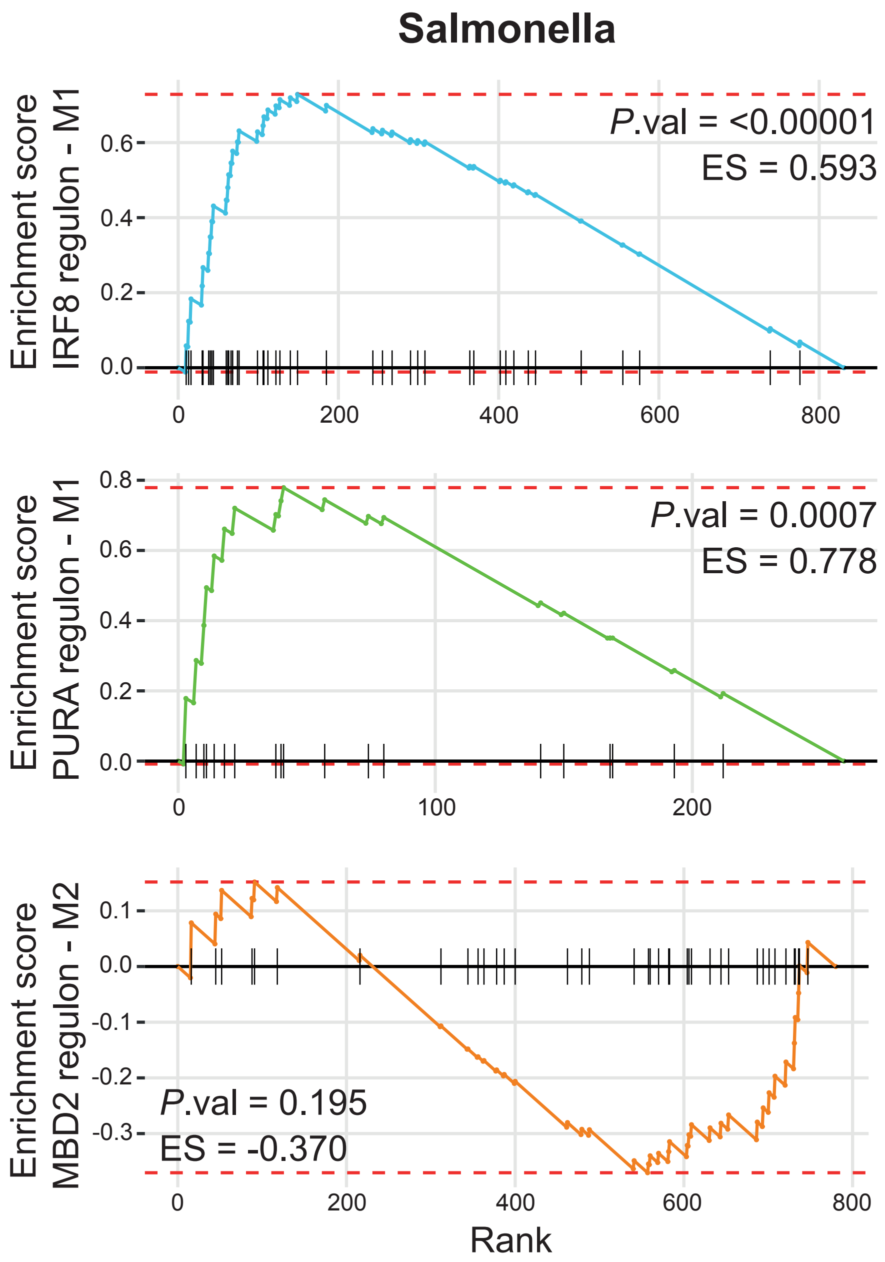


**<u>Supplementary Figure S18.</u> Gene set enrichment analysis (GSEA) identifies M1 signature in the IRF8 regulon.** GSEA of genes included in the IRF8 (top, blue), PURA (middle, green), and MBD2 (bottom, orange) regulons from the union of the naive and infected macrophage eGRNs, that were differentially expressed (*P*adj < 0.05) in salmonella infected macrophages versus control. Genes were ranked based on log2 fold-change (x-axis). Enrichment score is depicted as colored lines (y-axis), and vertical black bars below indicate the position of M1 associated genes for IRF8 and PURA, and M2 associated genes for MBD2. ES = normalized enrichment score, *P*.val = *P*-value.

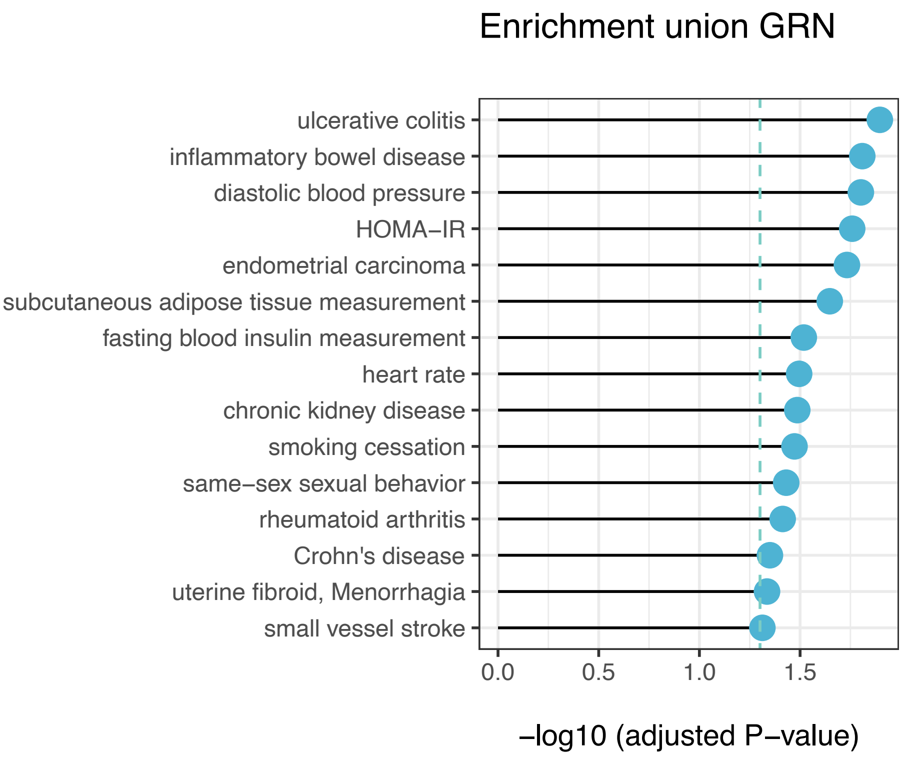


**Supplementary Figure S19:** Lollipop plot of LDSC enrichment P-values. GWAS traits that attained a nominal P-value < 0.05 for enrichment of heritability in the enhancer regions connected either NFkB1/2, RELB, or IRF8 in the union of the three macrophage eGRNs are displayed.

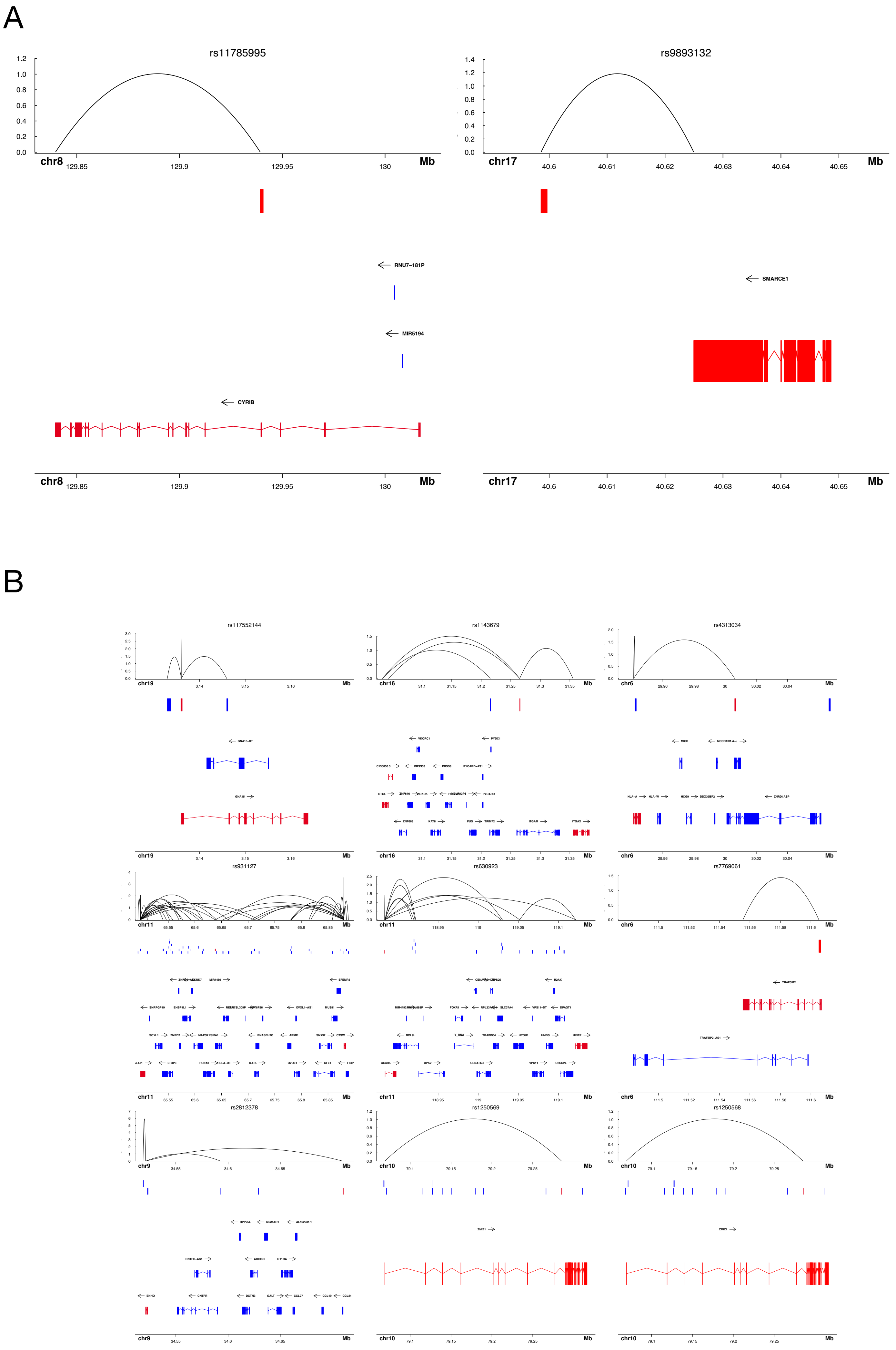
A
rs11785995
rs9893132
chr8
chr17
Mb
RNU7−181P
MIR5194
CYRIB
SMARCE1
B
rs117552144
rs1143679
rs4313034
rs931127
rs630923
rs7769061
rs2812378
rs1250569
rs1250568

C

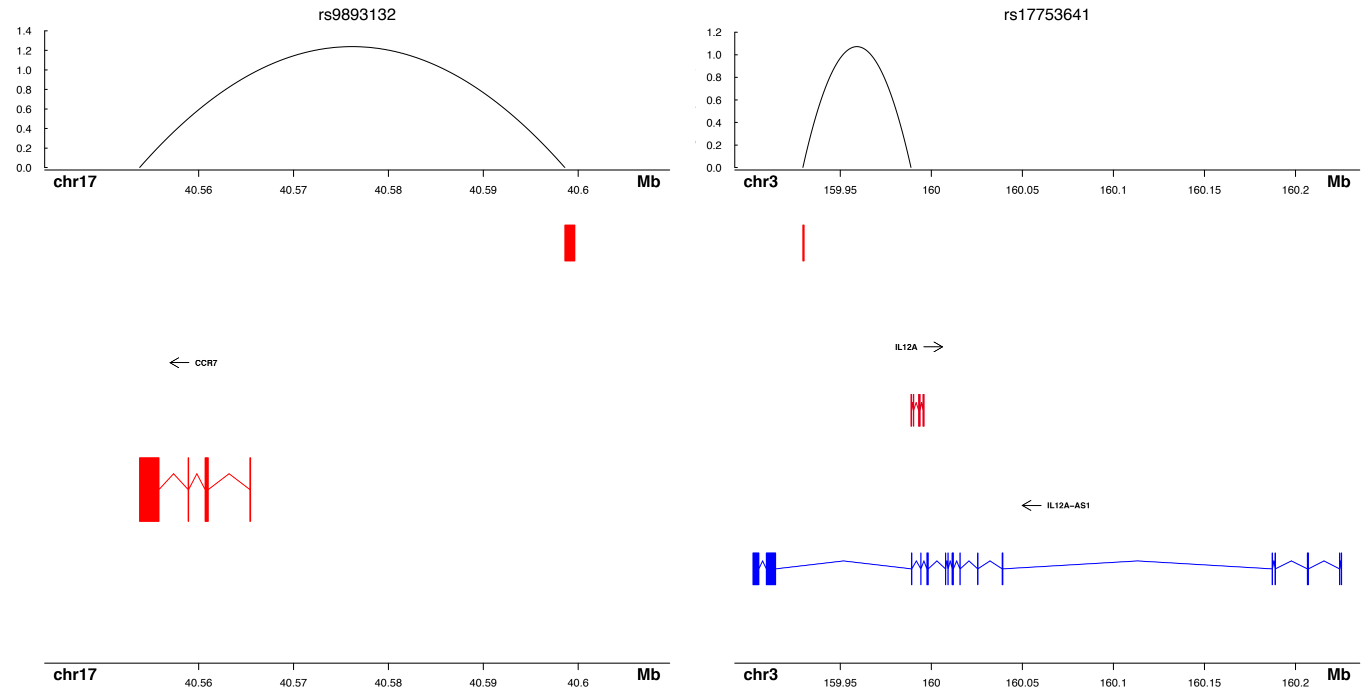


**Supplementary Figure S20**: The genomic context of all 11 fine-mapped variants in ATAC-seq peaks are shown in the naive (**A**), primed (**B**), and infected (**C**) macrophage eGRNs. The context includes gene tracks, other peaks present in the infected macrophage eGRN (blue boxes), and peak-gene links (arcs). Genes targeted by the peak overlapping with the SNP (red box) are colored in red.
